# Activation and regulation of a p38-MAPK by its downstream MAPKAP kinase through feedback phosphorylation and LLPS-driven condensate formation

**DOI:** 10.1101/2024.08.01.606155

**Authors:** Pranita Ray, Chakshu Mangal, T Pallavi Rao, Mintu Nandi, Afreen Haque, Vibha Viswanath, Samrat Mitra, Swasti Raychaudhuri, Anindya Ghosh-Roy, Smarajit Polley

**Affiliations:** Department of Biological Sciences, Bose Institute, Sector V, Bidhan Nagar, Kolkata 700091, India; Department of Cellular and Molecular Neuroscience, BRIC-National Brain Research Centre, Manesar 122052, India; CSIR-Centre for Cellular and Molecular Biology, Uppal Road, Hyderabad 500007, India; Academy of Scientific and Innovative Research (AcSIR), Ghaziabad-201002, India; Department of Chemistry, Indian Institute of Engineering Science and Technology, Shibpur, Howrah 711103, India

## Abstract

MAP kinases (MAPKs) represent a class of evolutionarily conserved stress and extracellular stimuli responsive signaling molecules. p38 group of MAPKs known to be activated by dual-specific upstream MAPK kinases and also by autophosphorylation. They activate MAPK activated protein kinases (MAPKAPKs) in a context dependent manner by specific phosphorylation, and together, they play crucial biological roles. One such pair is PMK3, p38α-MAPK and its cognate MAPKAPK, MAK2, downstream of DLK1 (MAPK kinase kinase) and MKK4 (MAPK kinase) in *C. elegans*. They are implicated in axonal regeneration, degeneration and synaptic pruning in response to neuronal injury. Here, we report that PMK3 and MAK2 engage in a feedback relationship, leading to phosphorylation-mediated activation of both kinases. Interaction of PMK3 with MAK2 through the canonical docking site leads to negligible *de novo* autophosphorylation of PMK3. Phosphorylation of both Thr and Tyr residues on the TxY-motif necessary for the full activation of PMK3 requires catalytic activity of MAK2 as confirmed by mass spectrometry. Distribution of phosphorylation sites on either kinase when incubated together, and presence of long intrinsically disordered regions in each kinase indicate that PMK3:MAK2 complex is conformationally plastic in nature. MAK2 increases bioavailability of aggregation-prone PMK3 by virtue of its ability to form LLPS-driven condensates *in vitro,* wherein they retain enzymatic activities and phosphorylate each other. Furthermore, experiments with transgenic *C. elegans* reveal that MAK2 controls stability of PMK3 and localization of PMK3 puncta in touch neuron. Our observations offer an unreported activation mechanism of a p38-MAPK by its downstream MAPKAPK.

**Significance Statement:** MAPK kinases are evolutionarily conserved, and are key players in stress response, cell survival, differentiation, metabolic processes and neuronal response to injury. MAPK pathways are primarily activated through unidirectional flow of phosphorylation-signal MAP3K to MAP2K to MAPK, and in some cases to a downstream MAPKAPK. We found that a *C. elegans* MAPK (PMK3) can also be activated by its downstream MAPKAPK (MAK2) through feedback phosphorylation that ensures robust activation of PMK3 by MAK2 without requiring the MAP2K. Furthermore, MAK2 increases bioavailability of activation-competent and active PMK3 by preventing its aggregation through LLPS-driven condensate formation *in vitro* and in the worm neuron. This feedback relationship might ensure rapid activation of such MAPK pathways in response to nervous system injury or stress.

## Introduction

Protein kinases are implicated in almost, if not all, signaling events in eukaryotes (1). Mitogen activated protein kinases (MAPKs) are activated by a variety of extracellular stimuli that regulate a plethora of cellular processes including but not limited to gene expression, cell cycle, cell survival, infection, inflammation, apoptosis, differentiation and metabolism (2–6). They are one of the most ancient signal transduction molecules found throughout evolution. Multiple MAPK pathways are found to operate in concert in eukaryotes. A canonical MAPK module consists of two upstream kinases, MAPK kinase (MAPKK or MAP2K) and MAPKK kinase (MAPKKK or MAP3K)(7). In response to extracellular stimuli, a MAP3K activates its cognate MAP2K that in turn activates the downstream MAPK – making it a three-kinase signaling module where the signal transmits from the MAP3K to MAPK via the MAP2K. Activation of p38α by autophosphorylation has also been documented (8) apart from the canonical mode of MAPK activation by MAP2K, suggesting multiple modes of MAPK-activation in play. MAPKs are Ser/Thr kinases that phosphorylate different kinds of protein substrates in cells including transcription factors and specific protein kinases called the MAPK activated protein kinases (MAPKAPKs) (9).

p38 group of MAPKs are perhaps the most widely implicated and most well-studied pathways among the four distinct subgroups of the MAPK family (2, 10). In mammals, there are four isoforms of p38: α, β, γ and δ. Among these p38α is the most well studied one. Among many substrates of p38α, MAPKAPK2 or MK2 have been implicated in many cellular processes that phosphorylate substrates including small heat shock protein 27 (Hsp27), cAMP response element-binding protein (CREB), lymphocyte-specific protein 1 (LSP1), transcription factor ATF1 etc. (11). p38 MAPKs are present both in the cytoplasm and nuclei of cells and are sequestered mostly in the nuclei in certain stress conditions. Nucleocytoplasmic shuttling of the p38 MAPKs is also regulated by their association with the MAPKAPKs (12).

*C. elegans* genome consists of three genes, *pmk1, pmk2* and *pmk3* that code for three p38 MAPK isoforms: PMK1, PMK2 and PMK3, respectively. These three genes constitute an operon with a polarity from *pmk-1* to *pmk-3*, and are expressed under the influence of a single promoter (13). Among these three p38 MAPK isoforms, PMK3 has been implicated in axonal regeneration acting as the MAPK downstream of MAP2K, MKK4 that is activated by the MAP3K, DLK1 in both *C. elegans* and other organisms (14–17) This signaling mechanism is also implicated in synaptic maturation in multiple organisms, which is under the tight regulation of E3 ubiquitin ligase (18–20). In the axonal regeneration pathway involving DLK1, the terminal kinase however is not the MAPK but a cognate MAPKAPK of PMK3 known as MAK2 akin to human MK2 or MK3 (Fig. 1A). MAK2 is involved in the maintenance of the neuronal synapses and axon morphology. It has also been shown that MAK2 regulates touch neuron axon regeneration by modulating *cebp-1* mRNA stability in response to axon injury in nematode (21). A search for the protein-protein interaction networks related to PMK3 and MAK2 on the STRING database revealed strong interactions between the two proteins (Fig. 1B), both in the full STRING network as well as physical subnetwork (22). We asked the question, whether activation, regulation and mutual relationship of p38α and its MAPKAPK established in mammalian systems also holds true in the PMK3:MAK2 system, which is involved in multiple cellular responses such as axon regeneration (23), degeneration (24), and synaptic pruning (25) following neuronal injury. We used the nematode specific counterparts of MAPK (PMK3) and MAPKAPK (MAK2) as our model systems. Our biochemical, computational, mass spectrometric and organismal analyses revealed that PMK3 participates in a hitherto unknown feedback relationship with MAK2. PMK3 activates its downstream kinase MAK2 that leads to the double-phosphorylation of the activating TxY motif in PMK3 leading to its full activation. We also found that, MAK2 but not PMK3 forms condensates through liquid-liquid phase separation (LLPS) *in vitro* and prevents aggregation of PMK3. Presence of MAK2 increases level of PMK3 in neurons under stress conditions in *C. elegans*. Further, the feedback-phosphorylation relationship remains functional in the condensate state. These observations point towards an unknown regulatory aspect of a MAP kinase (PMK3), wherein it is regulated by its cognate downstream kinase (MAK2), not only by means of its catalytic activity to chemically modify (activating phosphorylation) but also through change of physico-chemical microenvironment. Overall, our study revealed a new mechanistic understanding of the regulation of the injury sensing MAPK, PMK3 via its downstream kinase MAPKAPK, MAK2.

**Figure 1:**
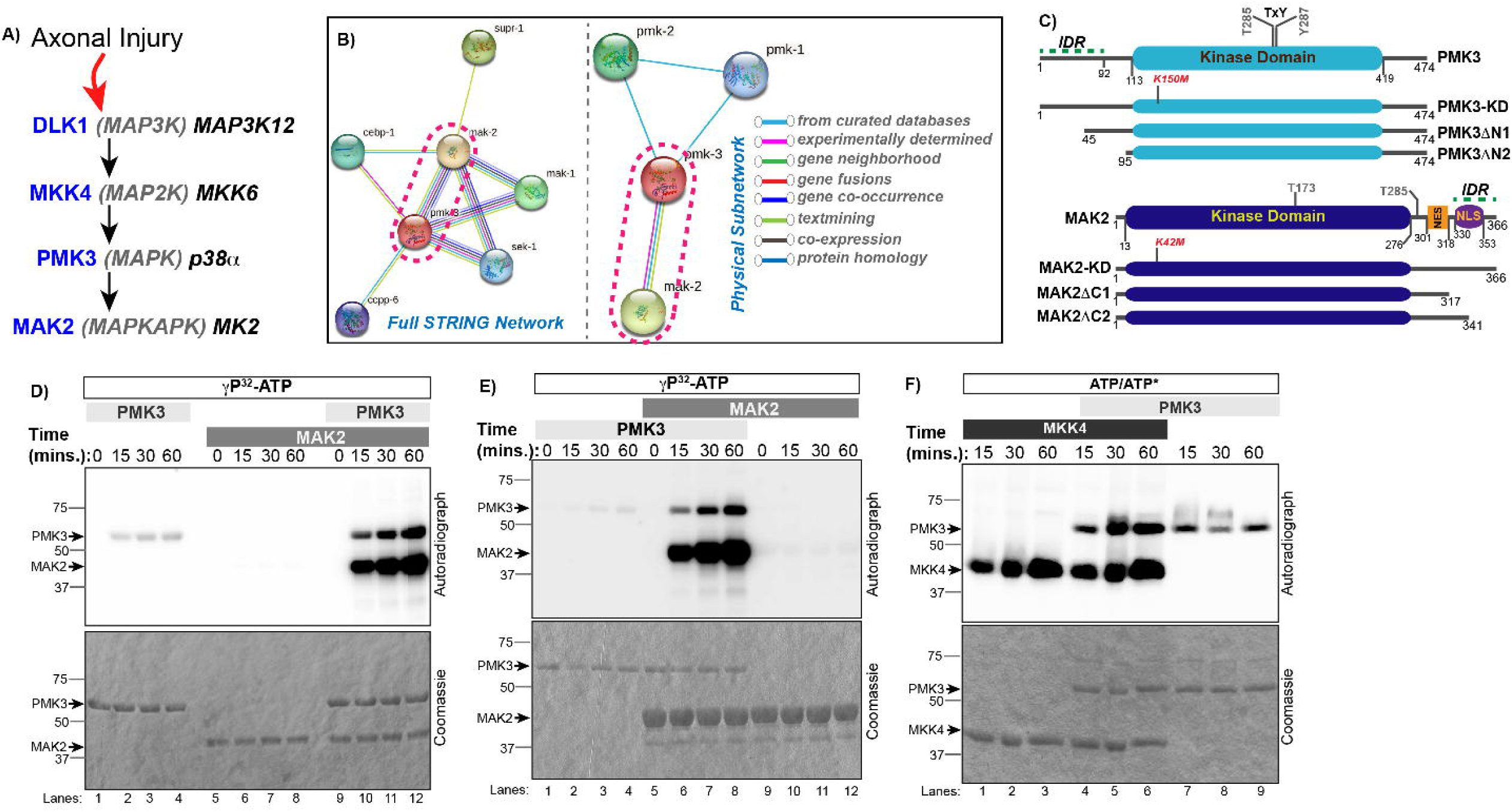
*In vitro* reconstruction of a MAPK-MAPKAPK module involved in axonal regeneration in *C. elegans.* **(A)** Schematic representation of the four-kinase DLK-1 MAPK signaling pathway in *C. elegans* (blue) along with their human counterparts (dark). PMK3 (MAPK) and MAK2 (MAPKAPK) act as the two terminal kinases in this module. **(B)** Analysis of the protein-protein interaction network between PMK3 and MAK2 using the STRING database. Full STRING network as well as physical interaction subnetwork are shown. **(C)** Top: Domain architecture of PMK3 depicting the kinase domain (Cyan) and the TxY motif within the activation loop. Lower panel: Domain architecture of MAK2 depicting the kinase domain (Blue), the nuclear export signal (NES, Orange) and the nuclear localization signal (NLS, purple). The domain borders have been numbered. For each kinase, location of the ATP-anchoring invariant lysine residues and predicted IDR regions are marked. Design of deletion constructs used in this study are also depicted. **(D-F)** *In vitro* radioactive phosphorylation assays with recombinant PMK3 (MAPK), MAK2 (MAPKAPK) and MKK4 (MAP2K). (D) *De novo* phosphorylation of PMK3 and MAK2 were monitored either alone or in presence of each other. (E) Same as (A) at a different relative stoichiometry to check reliance on the relative stoichiometry of the two constituent kinases. (F) Similar assay was performed to monitor directionality of phosphorylation signal transmission between PMK3 and its upstream kinase MKK4.

## Results

### In vitro reconstitution of a MAPK-MAPKAPK module implicated in axonal regeneration in C. elegans

PMK3 (Uniprot id: O44514, M.W. 54.9kDa) is a p38 MAPK in *C. elegans*. It participates in the DLK1-dependent signaling cascade in axonal regeneration pathway downstream of the MAP2K, MKK4 (KEGG pathway cel04361) (21, 23). PMK3 phosphorylates a downstream kinase MAK2 (Uniprot id: Q965G5, M.W. 41.5kDa), a MAP kinase activated protein kinase (MAPKAPK). Involvement of MAK2 in this cascade turns the canonical three-kinase MAPK-module into a four-kinase module (Fig. 1A). Schematic representation of the domain architectures and different constructs used in this study are shown in Fig. 1C. PMK3 shares highest sequence similarity with p38α among the human p38-isoforms (Fig. S1A), while MAK2 shares more than 50% identity with two human MAPKAPKs, MK2 and MK3 (Fig. S1B). To understand the mutual relationship between these two terminal kinases in further detail, we purified PMK3 and MAK2 recombinantly from *E. coli,* that were active in an *in vitro* radioactive kinase assay towards a generic substrate, myelin basic protein (MBP) (26) (Fig. S1C).

We employed *in vitro* kinase assay using γ-^32^P-ATP to check phosphorylation of MAK2 by PMK3. Purified PMK3 and MAK2, mixed at different ratios, were incubated for different time intervals at 25°C and *de novo* phosphorylation and autophosphorylation were monitored by autoradiography. It was observed that, both PMK3 and MAK2 were heavily phosphorylated in a time dependent manner when incubated together (lanes 9-12), but did not show appreciable autophosphorylation when incubated alone in the same assay conditions (lanes 1-8) (Fig. 1D). Enhancement of phosphorylation of MAK2 in presence of PMK3 was expected. However, profound increase in phosphorylation of PMK3 in presence of MAK2 may result from the direct catalytic activity of MAK2 or by virtue of physical interaction that leads to autophosphorylation of PMK3 (27, 28). This interdependence was independent of the relative stoichiometry of the two kinases as similar observations were made at two different molar ratios of the kinases used in Fig 1D (2μM of each) and 1E (1μM PMK3 and 6μM MAK2).

It is known that, active MKK4 is able to activate PMK3 by phosphorylation (19), similar to human p38α activation by MKK6 (29–31). PMK3 did not enhance *de novo* phosphorylation of its upstream kinase MKK4, in a similar assay involving PMK3 and MKK4, where MKK4 efficiently trans-phosphorylated PMK3 as expected (Fig. 1F).

### PMK3 and MAK2 engage in a feedback phosphorylation loop

Next, we wanted to further analyze the enhancement of phosphorylation of PMK3 and MAK2 in presence of each other (Fig. 2A). We took advantage of a mutant PMK3, where the invariant ATP binding Lys, K150 was replaced by Met, giving rise to K150M mutant or kinase dead version of PMK3 (PMK3-KD). This ATP-anchoring Lys, conserved in all eukaryotic protein kinases, helps position ATP in the kinase, resulting in a productive phospho-transfer reaction. Mutation of this Lys leads to abrogation of kinase activity; such mutants, in essence, are either kinase dead or retain minimal activity, if any. In a radioactive *in vitro* phosphorylation assay using PMK3-KD and MAK2, phosphorylation signal for PMK3-KD in the presence or absence of MAK2 (Fig. 2A, lanes 1-8), was hardly detectable. MAK2, in presence or absence of PMK3-KD, was faintly phosphorylated (Fig. 2A, lanes 5-12), but this phosphorylated form of MAK2 failed to enhance phosphorylation of PMK3-KD. Next, we used PMK3-activated phospho-MAK2 (P-MAK2) and monitored phosphorylation of PMK3-KD and P-MAK2 using the same experimental scheme. Phosphorylation of PMK3-KD in absence of P-MAK2 (i.e., auto-) was barely detectable, while significant phosphorylation of PMK3-KD was observed in presence of P-MAK2 (Fig. 2B, compare lanes 1-4 and 5-8). PMK3-activated P-MAK2, on the other hand, was capable of high level of self-phosphorylation as well (Fig. 2B, lanes 9-12). We also created a kinase dead version of MAK2 (K42M mutant, MAK2-KD) and found that, while it failed to undergo any self-phosphorylation (Fig. 2C, lanes 9-12), it was mildly trans-phosphorylated by PMK3 that showed very faint phosphorylation signal in these reactions (Fig. 2C, lanes 5-8). In comparison to MAK2-KD, MAK2 underwent significantly higher-level phosphorylation in presence of PMK3 (Fig. 2D). Likewise, PMK3 displayed much more efficient *de novo* phosphorylation signal in presence of MAK2 than MAK2-KD (Fig. 2D).

**Figure 2:**
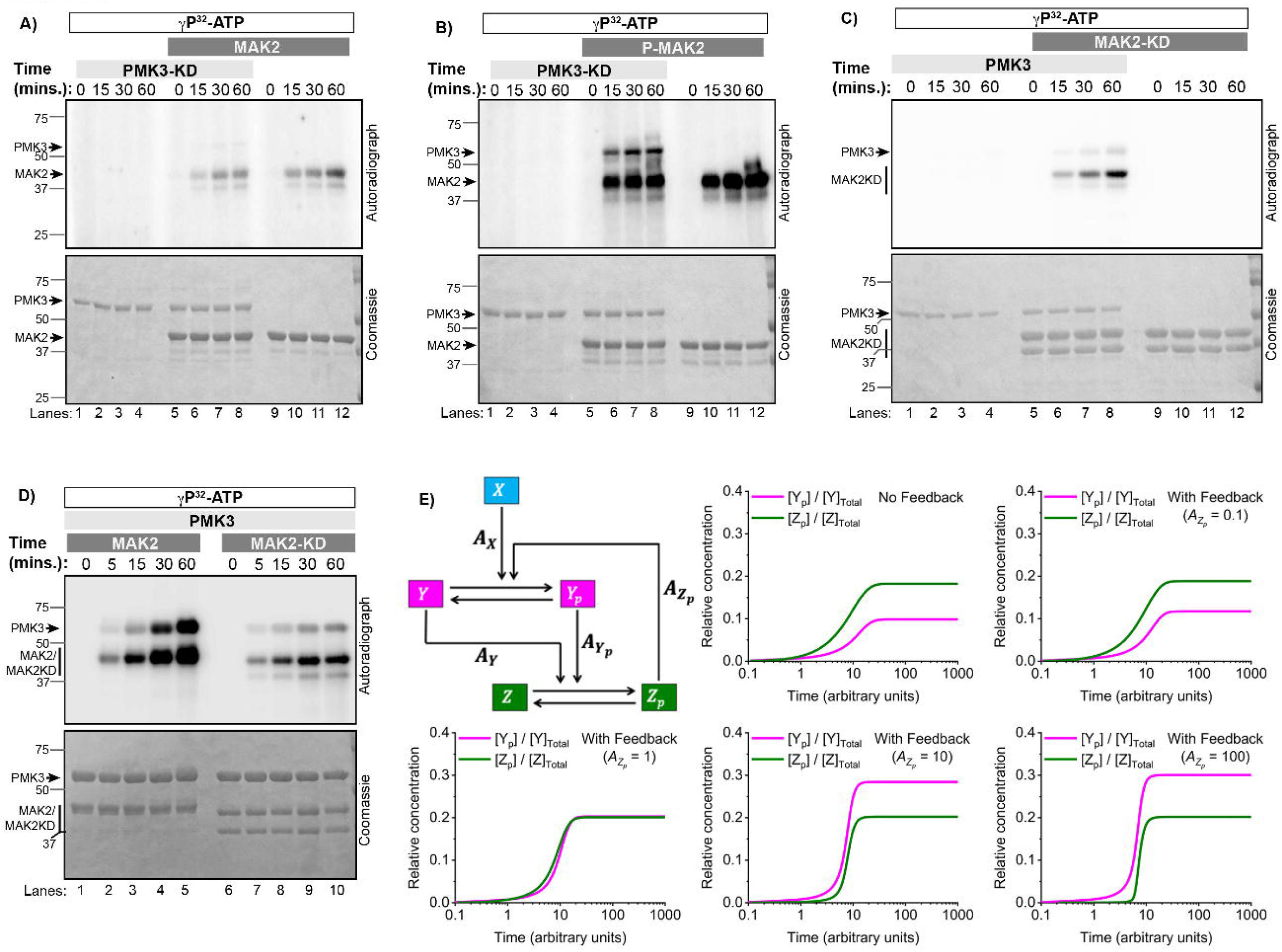
PMK3 engages in feedback phosphorylation with the downstream Kinase MAK2, but not the upstream kinase MKK4. **(A)** *de novo* phosphorylation status of a kinase dead PMK3 mutant (PMK3 K150M or PMK3-KD) and wild type MAK2 were monitored in an *in vitro* radioactive kinase assay at different time intervals. **(B)** Similar assay was performed with PMK3-KD and activated MAK2 (P-MAK2, pre-phosphorylated by PMK3) and their *de novo* phosphorylation status were monitored as a function of time. **(C)** Same as (A), except a kinase dead mutant of MAK2 (MAK2 K42M or MAK2-KD) and wild type PMK3 was used in this case. **(D)** *De novo* phosphorylation of wild type PMK3, MAK2 and MAK2-KD were monitored at different time points in two separate reactions where PMK3 was incubated with MAK2 or MAK2-KD. **(E)** Computational modelling of the effect of feedback loop on the overall phosphorylation kinetics of the two kinases. The temporal evolution of relative concentrations of Y_p_ (phosphorylated PMK3) and Z_p_ (phosphorylated MAK2) are presented with respect to total kinase amount of each kinase for different values of feedback strengths (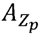), which is a dimensionless measure of feedback strength and sensitivity.

In order to understand the effect of such feedback relationship, we modeled the system as depicted in Fig. 2E; wherein X denotes MKK4 (MAP2K), Y denotes PMK3 (MAPK) and Z denotes MAK2 (MAPKAPK). Y_p_ and Z_p_ represent the phosphorylated versions of the respective kinases and are considered active. The model captures the effect of feedback on the phosphorylation kinetics and relative amounts of phosphorylated kinases (with respect to the total kinase amount) (Fig. 2E). The effect of feedback is accounted by considering a dimensionless parameter 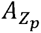, called feedback strength and sensitivity. Clearly, such feedback is advantageous to generate active pools of those kinases than without feedback.

### Mass spectrometric analysis of PMK3 and MAK2 phosphorylation

p38-MAPKs require phosphorylation of both T and Y in the TxY motif in their activation loops to usher maximum activity, while phosphorylation of only T show much reduced activity and that of only Y renders the kinase inactive (32). In the previous sections, we observed that, phosphorylation on both PMK3 and MAK2 increased many folds when they were both present in the reaction compared to the lone-kinases. However, identities of phosphosites are unknown. We performed high resolution LC-MS/MS experiments to identify phosphorylation sites after incubating PMK3, MAK2, and a mixture of PMK3 and MAK2 with ATP. We considered only those phosphorylated residues with probability of phosphorylation greater than 0.85 and that were detected at least twice (MS/MS count ≥2). List of those sites are presented in Table 1; for clarity the phosphosite-identification data are collated in Venn diagrams (Fig. 3A) and in individual cartoon representation of PMK3 and MAK2 (Fig. S2A and B, respectively). Evidently, more phosphorylated residues were identified in both PMK3 and MAK2 when they were incubated together, than individually with ATP. Most relevant phosphorylation event in the context of PMK3 activation is double phosphorylation of T285 and Y287 (pT285pY287). Singly phosphorylated peptides bearing pT285 or pY287 were detected in PMK3 alone treated with ATP, without the detection of any peptide containing pT285pY287, although peptides containing pT282pY287 or pS283pY287 were successfully detected in the same sample (Fig. 3A and B). No singly phosphorylated pT285 bearing PMK3 peptide was detected when incubated with ATP in presence of MAK2 although peptides bearing pY287 were observed (Fig. 3A and B). PMK3-peptides containing MAPK-activating pT285pY287 were detected exclusively in presence of MAK2 (Fig. 3B and C, S3C) along with multiple other doubly phosphorylated sites including those observed in the PMK3-only sample. Distribution of phosphorylation sites in the activation segments of PMK3 (Fig. 3D) and in MAK2 (Fig. S2D) are also shown. We also found, using a context-independent phospho-tyrosine specific antibody, that a small fraction of PMK3 was already tyrosine-phosphorylated that enhanced significantly when treated with ATP (Fig. 3E, lanes 1-4) and treatment with MAK2 further enhanced it (Fig. 3E, lanes 5-8). MAK2, on the other hand, was tyrosine-phosphorylated to a very small extent by PMK3 (Fig. 3E, lanes 5-8) while that on MAK2 in absence of PMK3 with or without ATP was barely detectable (Fig. 3E, lanes 9-12).

**Figure 3:**
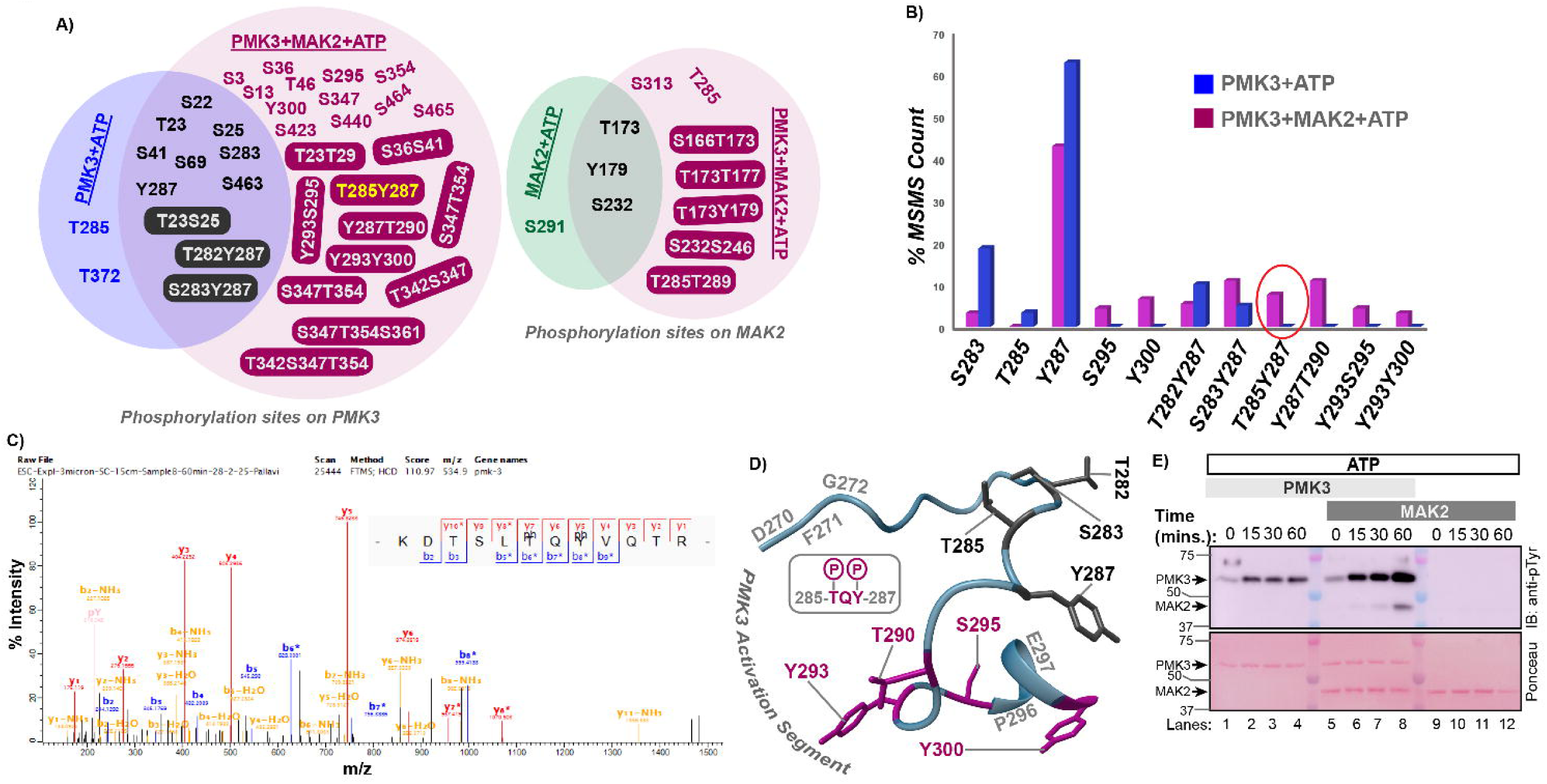
Identification of phosphorylation sites on PMK3 and MAK2. **(A)** Venn diagrams showing identities of singly and doubly-phosphorylated residues in PMK3 and MAK2 identified by LC-MS/MS analyses when incubated with ATP either individually or together. PMK3-residues phosphorylated in PMK3-only and with MAK2 samples are shown in blue and purple, respectively while residues common in both samples are depicted in dark letters (left panel). MAK2-residues phosphorylated in MAK2-only and with PMK3 samples are shown in green and purple, respectively and those common in both samples are depicted in dark letters (right panel). In each case, doubly-phosphorylated entities are shaded with respective colors. **(B)** Percentage occurrence of individual PMK3-phosphosite in presence (purple) and absence (blue) of MAK2 are compared. Percentage occurrence of phosphorylation for each phosphorylation site in PMK3 is calculated from MS/MS counts of each phospho-peptide in total phospho-peptide pool. Only those phosphorylation sites are considered having probability > 0.85 and MS/MS counts ≥2. **(C)** MS/MS spectrum of a PMK3-peptide containing doubly phosphorylated TxY motif (-KDTSLpTQpYVQTR-) is shown. **(D)** Ribbon representation of the activation segment of PMK3 from DFG- to APE-motif. Phosphorylation sites identified in this study are shown in sticks; wherein purple color depicts residues phosphorylated only in presence of MAK2 while those in dark gray are found both in presence and absence of MAK2. **(E)** PMK3 and MAK2 were incubated with non-radioactive ‘cold’ ATP and subjected to immunoblot analyses using a context-independent anti p-tyrosine antibody. The blot is a representative of three independent experiments.

In order to look at the spatial distribution of phosphorylation sites in three dimensions, two different structural models of the PMK3:MAK2 complex were generated using AlphaFold-predicted structures (33), due to the lack of any experimentally derived structure for either kinase or their complex. Firstly, we superimposed AlphaFold-derived structures of PMK3 and MAK2 on to the experimentally derived structure of the human p38α:MK2 complex (pdb id: 2oza), and also generated a model of the PMK3:MAK2 complex using AlphaFold2. When phosphorylation sites on both PMK3 and MAK2 were highlighted in these two structural models of the said complex, it became clear that it’s unlikely for either kinase to access all the phosphorylation sites on the other kinase from either pose (Fig. S2E and F, left and right panels). Neither could these kinases autophosphorylate themselves at sites far from the respective active sites. We speculate that PMK3:MAK2 complex is dynamic in nature giving rise to an ensemble of conformations, and the experimentally derived pose of the human p38α:MK2 complex or AlphaFold2-derived PMK3:MAK2 complex are insufficient to capture the dynamicity and/or plasticity of the said complex.

### Sequence and structural analysis of PMK3 and MAK2

To glean into the complex between PMK3 and MAK2, we used the above-mentioned superimposed-complex structure (pdb id: 2oza) and analyzed the CD and ED domains on PMK3 that are major contributors in recognizing the MAPKAPKs (34, 35). Residues in the ED domain are largely conserved with the exception of N159 of p38α changed to another polar residue T261 in PMK3, and C162 of p38α changed to S264 in PMK3, that changes −SH to −OH (Fig. S3A). In the CD domain, however, changes are more drastic; acidic D313 of p38α changed to hydrophobic L424 in PMK3 and polar, phosphorylatable Y131 of p38α changed to hydrophobic, phospho-ablative F422 in PMK3 (Fig. S3B). When we turned our attention to the p38-docking site (R/K–R/K–X_10_–R/K–R/K–R/K–R/K–R/K), we realized that this region of MAK2 is not fully conserved compared to its human counterparts MK2 and MK3 (Fig. S3C). For example, 2^nd^ R/K (K374 of MK2) and last R/K (K389 of MK2) have been changed to S325 and G340, respectively in MAK2 (Fig. S3C). Such changes in the interacting residues among the partner kinases of human and nematode origins raise the possibility that the ensuing PMK3:MAK2 and p38α:MK2 complexes may differ in spatial arrangements as well as affinity.

We also noticed that PMK3 contains a long (~90aa) insert at the N-terminus before the kinase domain that is absent in the human p38 isoforms (Fig. S1A). On the other hand, MK2 and MK3 contain long N-terminal insertion before the beginning of the kinase domain that is absent in MK5 and MAK2 (Fig. S1B). MAK2, however contains a highly charged C-terminal region beyond the MAPK-docking sequence that is absent in MK2 or MK3 (Fig. S1B). Both the N-terminal extension in PMK3 and C-terminal extension in MAK2 are largely unstructured in AlphaFold2-predicted structures (Fig. 4A). Next, we analyzed those sequences using web-based tools, like MolPhase, FuzDrop, IUPred3, PONDR and FoldIndex, to gain further insights on their physico-chemical properties (36–39). For PMK3, pLDDT score was lowest at the N-terminus prior to the kinase domain and the same region was predicted to be intrinsically disordered by each of the predictive tools employed (Fig. 4A, left panel and S4D); the same was true for the MAK2 C-terminus beyond the kinase domain (Fig. 4A, right panel and S4E). Presence of intrinsically disordered regions (IDRs) bestow their parent proteins with multivalency of interaction (40, 41), and presence of long IDRs in both PMK3 and MAK2 may posit each other in ways to result in an ensemble of conformation making phosphorylation at different surfaces possible. Notably, MAPK-docking site on MAK2 resides in its C-terminal IDR (42).

**Figure 4:**
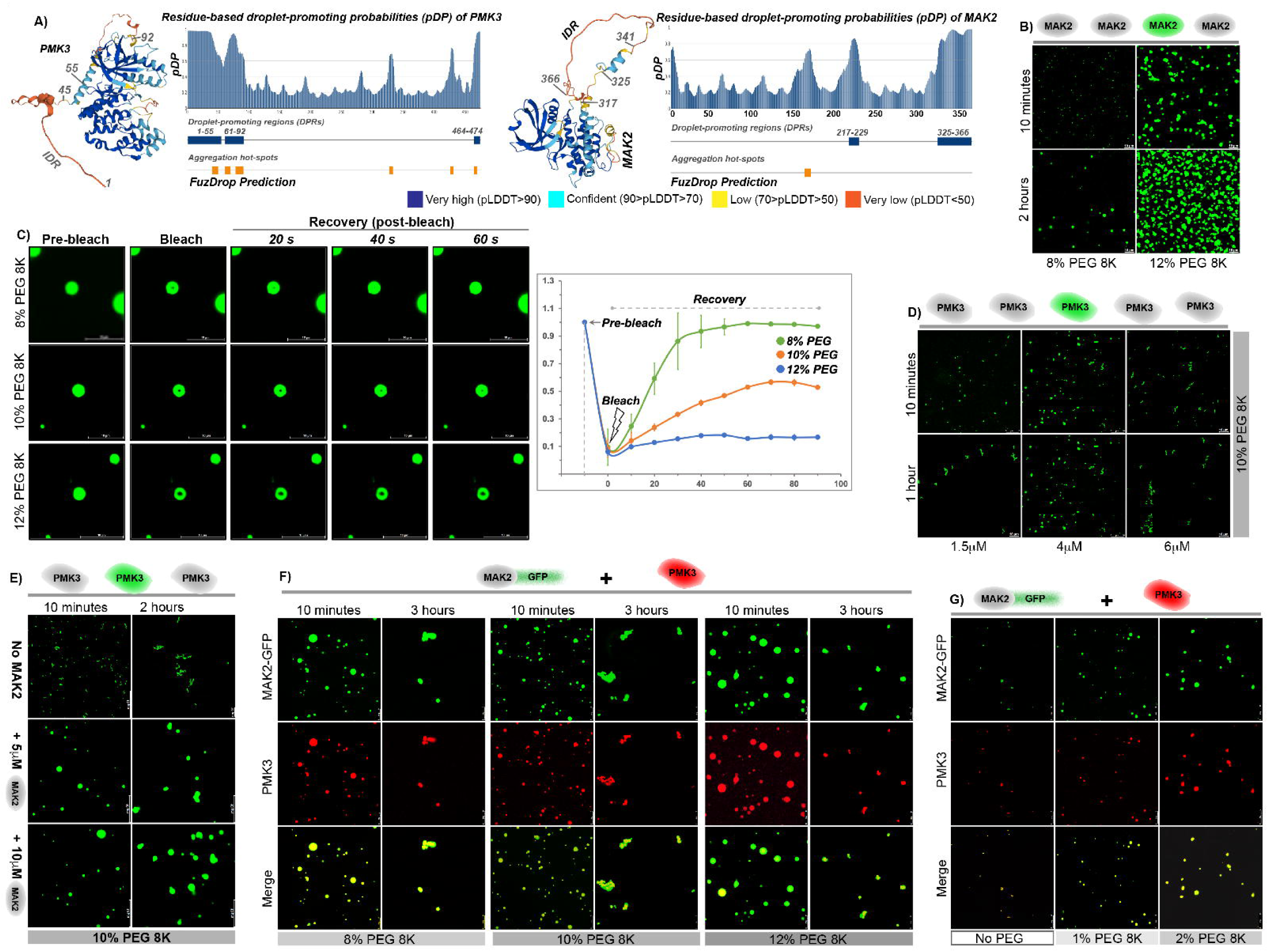
PMK3 and MAK2 harbor unstructured, disordered regions and propensity to undergo phase separation. **(A)** Left panel: FuzDrop prediction shows the droplet promoting regions (DPRs) along with the aggregation hotspots in the PMK3 sequence, AlphaFold predicted structure of PMK3 colored according to the pLDDT score (signifies confidence of prediction) is also shown. Right panel: Shows FuzDrop analysis of MAK2 and its AlphaFold predicted structure colored in the same manner as PMK3 structure. The number of aggregation hot spots in the MAK2 sequence is distinctly lower than those found in PMK3. Region spanning amino acids 325-341 contains the MAPK-docking site. Color coding of the pLDDT score is shown at the bottom. **(B)** Fluorescein labelled MAK2 (F-MAK2) was incubated with varying concentrations of PEG-8000 for different time periods and imaged under the microscope. MAK2 undergoes phase separation to form the round droplets under both the conditions. Images are representative of three independent experiments. Scale bar is 10μm. **(C)** FRAP analysis on the phase separated MAK2-GFP condensates at different PEG concentrations (left). Scale bar is 10μm.The fluorescence recovery in each case was followed up to 60 seconds post bleaching and the normalized fluorescence intensity was plotted against time (right). The data values are shown as mean±s.d. of normalized fluorescence intensity values from three independent measurements (n=3). **(D)** *In vitro* droplet formation assays with fluorescein labelled PMK3 show that PMK3 forms aggregates at different protein concentrations in the presence of 10% PEG-8K. Images are representative of three independent experiments. Scale bar is 10μm. **(E)** F-PMK3 was mixed with unlabeled MAK2 at two different concentrations in presence of 10% PEG and imaged at 10 minutes and 2 hours respectively. In presence of MAK2, PMK3 undergoes liquid-liquid phase separation to form globular droplets instead of aggregates. Images are representative of two independent experiments. Scale bar is 10μm. **(F)** MAK2 and PMK3 co-localize within the same phase separated droplets. MAK-GFP (green) and Alexa 594 labelled PMK3 (red) were mixed together in presence of varying concentrations of PEG-8K (8-12%) and imaged at two time points. Merged (yellow) images show the co-localization of the two kinases within a single droplet. At 3 hours, the droplets exhibit a tendency to form aggregates with an increase in PEG concentration. Images are representative of two independent experiments. Scale bar is 8μm. **(G)** In presence of MAK2-GFP, PMK3 undergoes LLPS even in the absence of any crowding agent or as low as 1% PEG-8K. Scale bar is 8μm.

### MAK2 undergoes liquid-liquid phase separation and prevents aggregation of PMK3

Presence of IDRs also gives rise to the possibility of formation of molecular condensates through liquid-liquid phase separation (LLPS). MolPhase scores of 0.995 and 0.939 for PMK3 and MAK2, respectively, prompted us to investigate this possibility. We found that Fluorescein-labeled MAK2 (F-MAK2) forms condensates through LLPS at PEG concentrations of 8% and above (Fig. 4B), and both the number and size of the condensates increased with the increase in PEG concentration. At 12% PEG, the condensates seemed to be transitioning from a liquid-like state to an arrested state. To further confirm that LLPS in MAK2 was not due to labeling of the protein with Fluorescein, we generated and purified a MAK2 construct containing GFP at its C-terminus (MAK2-GFP) that also underwent LLPS to form condensate in a similar range of PEG concentrations (Fig. S3F). Size of these condensates increased as protein concentration was increased from 6uM to 10uM or more (Fig. S3G). To assess the liquid like properties of the condensates, we performed FRAP experiments at different concentrations of PEG (43). It was observed that the fluorescence fully recovered within 60s of bleaching at 8% PEG, while it was gradually arrested as the PEG concentration was increased to 10% and 12% (Fig. 4C). We also found that LLPS of MAK2 was severely compromised at higher concentration of NaCl (Fig. S3H). It suggests that MAK2 LLPS is a reversible process, energetics of which is likely dominated by electrostatic interactions (44, 45).

Unlike MAK2, Fluorescein-labeled PMK3 (F-PMK3) formed aggregates instead of phase-separated droplets in the same conditions that lead to LLPS-mediated droplet formation in MAK2 (Fig. 4D). Furthermore, this aggregation of F-PMK3 could not be prevented by the chaperone HSP90 or a non-interacting non-specific protein, BSA (Fig. S3I). Of note, low MW fraction of PMK3 displayed very high activity towards MAK2 (lane 6), while large MW fraction of PMK3 did not show any detectable activity (lane 3), indicating that PMK3 present in large self-assembly was inactive (Fig. S3J).

Next, we sought to investigate if MAK2 could prevent the aggregation of PMK3 and lead to LLPS-mediated droplet formation in PMK3. To this end, we took a fixed concentration F-PMK3 and incubated it with 10% PEG 8K in absence or presence of increasing concentration of unlabeled MAK2, and monitored F-PMK3 droplet formation at different time interval. Interestingly, instead of aggregating, F-PMK3 formed droplets through LLPS in presence of MAK2 (Fig. 4E). We also observed that, Alexa Fluor 594-labeled PMK3 (AF-PMK3) and MAK2-GFP colocalized in the same droplets as early as 10 mins post incubation (Fig. 4F), when MAK2-GFP and AF-PMK3 were used in a similar assay. At 3 hours post incubation, droplets showed tendency to aggregate with increasing concentration of PEG (Fig. 4F). Interestingly, AF-PMK3 and MAK2-GFP complex formed droplets through LLPS at PEG concentration as low as 1-2% and very few droplets were seen even when no PEG was present (Fig. 4G). These assays strongly bolstered the idea that MAK2 specifically engages with PMK3 to prevent its aggregation and it’s not a non-specific bystander effect.

### MAK2 regulates stability and localization of PMK3 in neuron

Our findings suggested that PMK3 is prone to aggregation and presence of MAK2 may prevent it to a large extent through LLPS *in vitro*. We wanted to investigate whether this MAK2 dependent regulation of PMK3 holds true within *C. elegans* neuron. Transgenic expression of PMK3-TagRFP reporter in touch neuron showed that PMK3 protein majorly localized to the cell body (Fig. 5A “WT” panel; named WT because it is Wild-Type for *mak-2*; arrow pointing PLM cell soma) and its nuclear localization was obvious. However, the intensity of PMK3-TagRFP was dim in the cell body, and it was hardly visible in both anterior and posterior neurites of PLM neuron (Fig. 5A “WT” panel). In the deletion mutant for *mak-2*, the intensity of the PMK3-TagRFP reporter gets further dimmed within PLM soma (Fig. 5A “*mak-2(lf)*” panel). Next, we co-injected PMK3-TagRFP and TagBFP-MAK2 in 1:1 and 1:6 ratios for co-expressing them in touch neurons, named “*mak-2(+)*” and “*mak-2(++)*”, respectively. When MAK2 was co-expressed, the intensity of PMK3-TagRFP was elevated in the cell body (arrow, Fig. 5A “*mak-2(+)*” panel). Interestingly, PMK3-TagRFP was visible within the neurites in presence of MAK2 (arrowheads, Fig. A “*mak-2(+)*” panel). Average intensity of PMK3-TagRFP was significantly higher in *mak-2* overexpression backgrounds as compared to wildtype in both the cell body and neurites, although diffusible GFP intensity remained unchanged (Fig. 5B and 5C, Fig. S4A). Of note, PMK3 intensity was slightly higher in the 1:6 expression line as compared to the 1:1 line (Fig. 5B and 5C). Same trend was observed in both L4 (larval) and aged animals (Fig. S4B and C). This suggested that MAK2 helps increase the stability and thereby intensity of PMK3 in both cell body and neurites of PLM neuron. Similar phenomenon was also noticed in ALM neuron.

**Figure 5:**
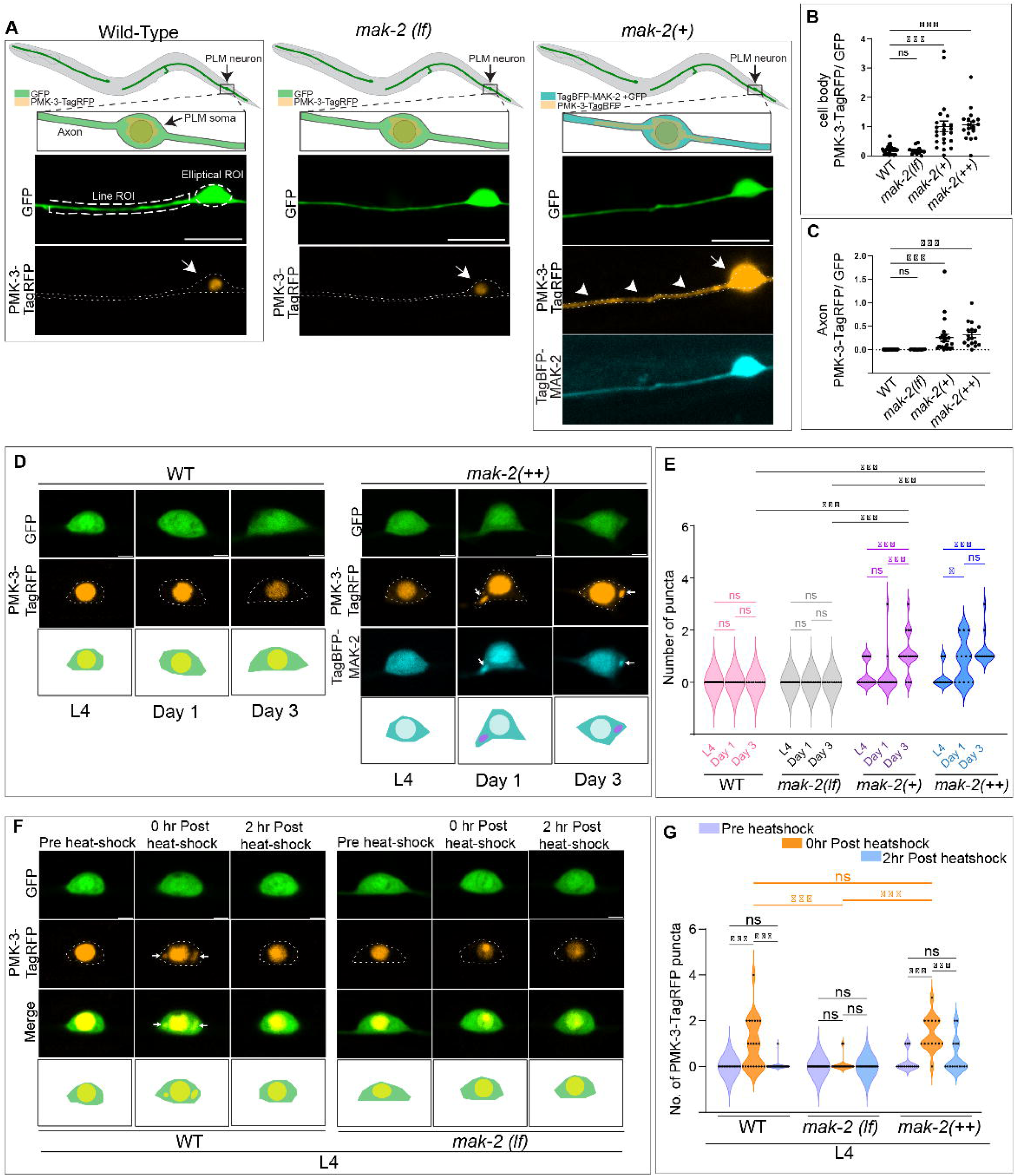
MAK2 regulates stability and localization of PMK3 in neuron. (A) The expression of P*mec4*-PMK3-TagRFP using a transgenic extra-chromosomal array (*shrEx522*) shows that PMK-3 localizes to the cell body (including nucleus) of posterior gentle-touch (PLM) neuron of C. elegans (“WT” panel, named WT because it is Wild-Type for *mak-2*), arrow pointing PLM cell soma). PMK3-TagRFP intensity gets dimmer in *mak-2*(ok2394) background. Co-expression of P*mec4-*PMK3-TagRFP and P*mec4-*TagBFP-MAK2 in a transgenic extra-chromosomal array (*shrEx526*) of 1:1 ratio, named “*mak-2*(+)”, shows that PMK3-TagRFP intensity is brighter both in the cell body (arrow, “*mak-2*(+)” panel) and axon (arrowheads, “mak-2(+)” panel) (Scale bar - 10 μm). (B) Scatter-plot showing the ratio of the mean intensity of PMK3-TagRFP vs diffusible GFP in region of interest (ROIs) in cell body (elliptical ROI-major axis=6 μm, minor axis=4 μm) in different genetic backgrounds. Increasing trend of mean-intensity ratio between PMK3-TagRFP and diffusible GFP in *mak-2* overexpression backgrounds as compared to wildtype in PLM cell body and axon (anterior neurite). (C) Scatter-plot showing the ratio of the mean intensity of PMK3-TagRFP vs diffusible GFP in region of interest (ROIs) (~ 60-μm line ROI of 5-pixel thickness) in PLM axon in different genetic backgrounds. (D) High-resolution images of the cell body region of PLM neuron in wild type and in a transgenic extrachromosomal array (*shrEx527*) co-expressing P*mec4-*PMK3-TagRFP and P*mec4-*TagBFP-MAK2 in 1:6 ratio, named “*mak-2*(++)” (Scale bar - 2 μm). (E) Quantification of number of PMK3-TagRFP/TagBFP-MAK2 puncta in PLM soma upon aging. The violin plot shows that the penetrance of puncta phenotype among animals increases with age. N=1, n~15 for each genotype. (F) The high-resolution images of the PLM cell body region for Pmec4-PMK3-TagRFP localization before and after heat shock. Upon heat-shock at 34deg for 2 hours, PMK3-TagRFP puncta can be seen in PLM soma. After recovery at 20 degrees, the puncta disappear. This puncta phenomenon is perturbed in *mak-2*(ok2394) (Scale bar - 2 micron). (G) Quantification of puncta number in L4 (larval) PLM soma pre- and post-heat shock and recovery. N=2, n~30 for each genotype except for mak-2(++) where N=1, n~15. Each dot in the graphs represent number of puncta per worm. Statistics: One-way Anova with Tukey’s multiple comparison test, P<0.001(***), <0.002(**), <0.033(*), <0.12(ns).

We further noticed that when TagBFP-MAK2 was expressed with PMK3-TagRFP, it resulted in infrequent puncta (condensed structures) formation in the PLM soma of L4 stage animals (Fig. 5D, “*mak-2*(++)” panel), although these puncta did not appear in transgenic line which exclusively expresses the PMK3-TagRFP reporter (Fig. 5D, “WT” panel).This observation was made in both 1:1 and 1:6 lines (arrows, Fig. 5D “*mak-2*(++)” panel, Fig. S4D). Also, the percentage of worms showing puncta phenotype increased with age (Day3 vs L4 data, Fig. 5E). This indicated that age related stress might be the factor that promotes these puncta formation, which requires MAK2.

We speculated that MAK2 could be helping in the formation of such condensed structures in response to stress to accommodate the aggregation-prone PMK3 (46, 47). In order to test this hypothesis, we used temperature as a stress factor and checked the effect of heat-shock on PMK3 in presence and absence of MAK2. Of note, heat-shock was shown to enhance LLPS conditions in other studies (48, 49). After heat shock at 34^0^C for two-hours, the condensed puncta of PMK3-TagRFP in cytoplasm of PLM soma increased significantly (Fig. 5F and 5G). These puncta were reversible as they disappear after animal was allowed to recover at 20^0^C for same time-period. Whereas in *mak-2* mutant background, such puncta formation was not seen for PMK3::TagRFP upon heat shock (Fig. 5F and 5G); however, PMK3::TagRFP intensity permanently decreased in those animals even after recovery at 20^0^C (Fig. 5F).

These results indicate that stress condition (like heat shock) causes aggregation of (possibly misfolded) PMK3 which can be reversed by endogenous MAK2 by preventing aggregation of PMK3 at favorable conditions. We speculate that PMK3 could be getting degraded by one of the degradation pathways in absence of MAK2. Similar results were obtained upon heat-shocking the animals at the Day1 and Day3 adult stage and with *mak-2(++)* line (Fig. 5G, Fig. S4E). Overall, these experiments demonstrate that MAK-2 regulates PMK-3 localization and stability *in vivo*.

### Effect of IDRs and phosphorylation on the feedback relationship of MAK2 and PMK3

It was evident from our biochemical analyses that MAK2 and PMK3 participate in a feedback phosphorylation (trans-) loop, whereas they undergo very little, if any, self-phosphorylation when incubated alone with ATP. To investigate how these self- vs. trans-phosphorylation modulate LLPS-mediated droplet formation and *vice-versa*, we designed three different reaction conditions, sets A, B and C as outlined in Fig. S5A. In set A, PMK3 was incubated alone in presence of 10% PEG 8K, where PMK3 aggregates, and then MAK2 was added. In this condition, only the fraction of PMK3 that is available in solution or condensate would serve as the kinase to phosphorylate MAK2 and as the substrate to be phosphorylated by MAK2. In set B, MAK2 was incubated alone in presence of 10% PEG 8K that is favorable for MAK2 condensate formation, and then PMK3 was added. In set C, PMK3 and MAK2 were incubated together in 10% PEG 8K, where the PMK3:MAK2 complex should form condensate. Both in sets B and C, larger fraction of PMK3 should be biochemically available to serve as the kinase or the substrate compared to set A. As expected, it was observed that in case of set A, phosphorylation signal for PMK3 was much less as not much of PMK3 was available in solution/condensate, and only a catalytic amount of PMK3 was available to phosphorylate MAK2 (Fig. 6A, left panel, lanes 1-3). In sets B and C, where PMK3:MAK2 complex was allowed to form condensates, both proteins underwent significant phosphorylation (Fig. 6A, left panel, lanes 4-9). We also confirmed that MAK2 was able to phosphorylate PMK3 on tyrosine(s), when both were present in the condensate (set C) (Fig. 6A, right panel) as was observed when both were present in solution (Fig. 3E). Next, we checked the effect of ATP-treatment on LLPS of MAK2-GFP and AF-PMK3 by incubating individual proteins along with 200μM of ATP at 8% PEG-concentration for different time intervals. They behaved similarly as to form droplets through LLPS for MAK2 and form aggregates for PMK3 when compared with the same protein in absence of ATP (Fig. S5B and C). Together these results inform us that autophosphorylation, if any, or in other words, mere presence of ATP is not sufficient to change PMK3’s tendency to aggregate or MAK2’s ability to from condensates through LLPS. MAK2:PMK3 complex underwent droplet formation through LLPS both in absence (Fig. S5D), as was already seen, and presence of ATP (Fig. 6B). Interestingly, we noticed that after prolonged incubation, the complex also showed tendency to aggregate in absence of ATP but that tendency to aggregate was not apparent in presence of ATP as the droplets remained rounded even after 3hrs post-incubation. These results show that MAK2 is able to form condensates and incorporate PMK3 into those condensates both in presence and absence of ATP, which might increase the bio-availability of PMK3 and activate it in presence of ATP.

**Figure 6:**
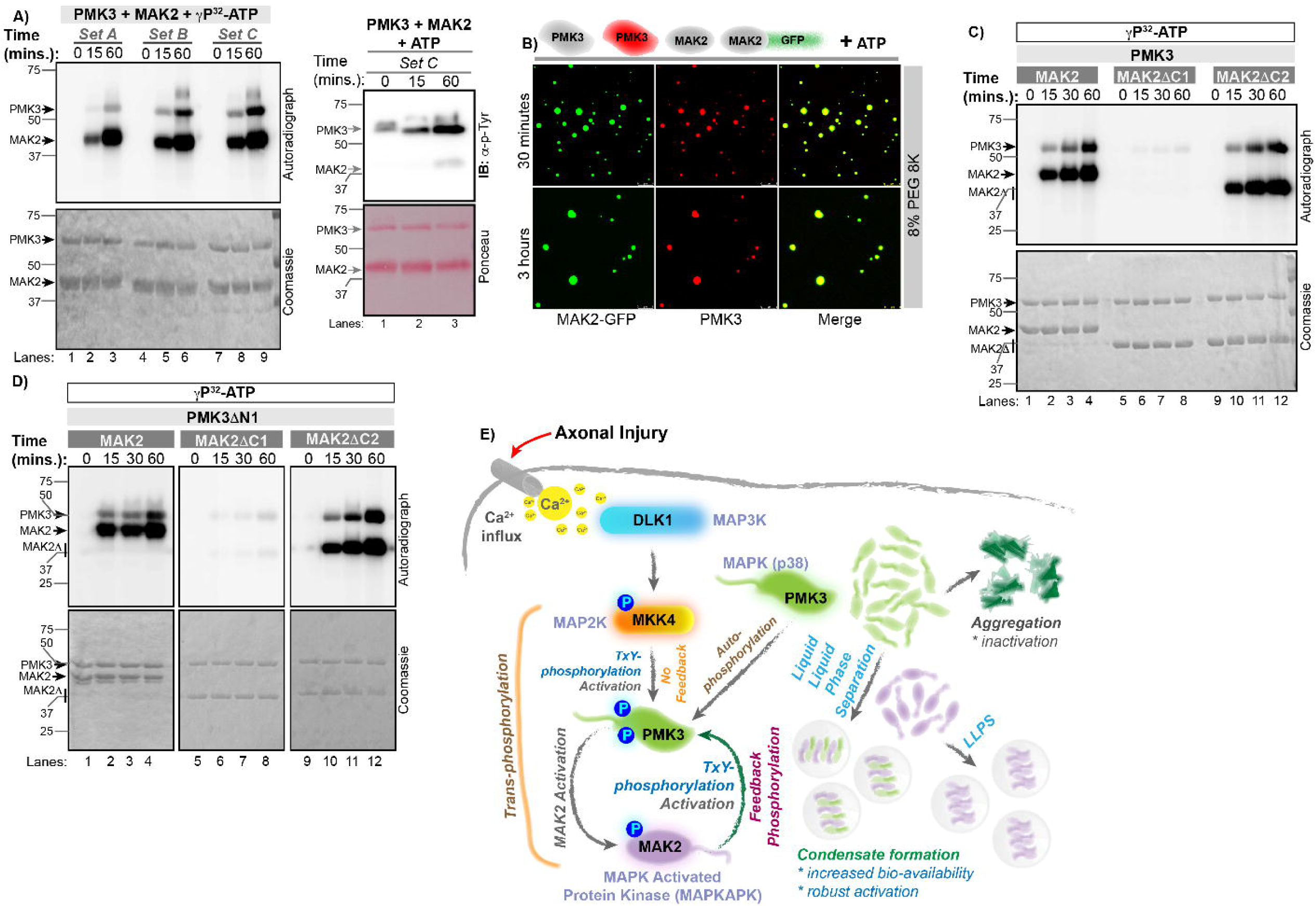
Effect of auto- and feedback phosphorylation on LLPS of the MAK2-PMK3 complex. **(A)** Left panel: PMK3 and MAK2 were incubated with radioactive ATP as per the reaction scheme shown in Figure S6, resolved by SDS-PAGE and *de novo* phosphorylation was monitored by autoradiography. Right panel: Reaction scheme C was performed with cold ATP and immunoblotting was performed using anti p-tyrosine antibodies to detect context-independent phosphorylation at the tyrosine residues of the two kinases. The blot is a representative of three independent experiments. **(B)** Effect of trans-phosphorylation on PMK3: MAK2 condensates. Scale bar is 8μm. In presence of ATP, the PMK3: MAK2 droplets maintained their globular look even at 3 hours post incubation. **(C)** *In vitro* radioactive kinase assay was performed using full length PMK3 incubated separately with full length MAK2 and two deletion constructs of MAK2, and *de novo* phosphorylation of each protein was monitored at different time points using autoradiography. **(D)** Similar assay, as described in *C* was performed using the same constructs of MAK2 with an N-terminal deletion construct of PMK3. **(E)** A graphical model depicting mutual relationship and signal transmission through PMK3 and MAK2. PMK3, a neuronal p38-MAPK is activated by its downstream MAPKAPK MAK2 without requiring the presence of canonical activator MAP2K, MKK4. MAK2 increases bioavailability of aggregation prone PMK3 by LLPS-driven condensate formation.

Next, we wanted to check the effect of the predicted intrinsically disordered regions (IDRs) in each kinase on their interdependence. In MAK2, the major IDR was predicted to be between residues 325-366 region; we created and purified two deletion constructs, MAK2ΔC1 and MAK2ΔC2 encompassing residues 1-317 and 1-341, respectively (Fig. 1C). MAK2ΔC2 behaved similarly as the full length MAK2 that displayed enhancement of PMK3 phosphorylation but MAK2ΔC1 was unable to do so, wherein only faint phosphorylation signal of PMK3 was visible (Fig. 6C). We also created two deletion constructs for PMK3, PMK3ΔN1 and PMK3ΔN2 encompassing residues 45-474 and 95-474, respectively as the predicted IDRs were located between residues 1-92 (Fig. 1C). Of note, we could successfully express and purify PMK3ΔN1 but not PMK3ΔN2. Similar observations were made for PMK3ΔN1 that displayed very little autophosphorylation, and full length MAK2 and MAK2ΔC2 efficiently phosphorylated it while MAK2ΔC1 couldn’t (Fig. 6D). Indeed, with gradual shortening of the IDR of MAK2 its ability to undergo LLPS went down and shorter constructs formed progressively smaller droplets compared to the full length MAK2 in similar assay conditions (Fig. S5E).

## Discussion

MAPK signaling modules have remained in the forefront of signal transduction research for many years. Many research articles employing biochemical, structural, genetics and other approaches helped the field to understand the importance of these kinases and detailed regulatory mechanisms (9, 50, 51). Studies of MAPK signaling in multicellular (mammalian) and unicellular (yeast) organisms show that the signal flows in a unidirectional path starting from a MAP3K that activates a cognate MAP2K to activate its downstream MAPK upon encountering extracellular stimuli. Basic structure and skeleton of this three-kinase module is preserved wherein MAP2Ks are dual specificity kinases required to phosphorylate the conserved TxY motif on the activation loop of the MAPKs, essential for the MAPKs to realize their full scale of activity (32). The three-kinase MAPK modules exhibit their roles either by directly phosphorylating effector-substrates (like transcription factors) or by activating a downstream kinase, MAPKAPK, by phosphorylation that ultimately carry out the phosphorylation of the effector-substrates (9). This makes the respective module a four-kinase MAPK module. Role of such MAPK-MAPKAPK duo are well established in the literature and have been implicated in mammalian biology, especially in the realm of immune signaling (52–54). Heterodimeric p38α:MK2 complex is known to orchestrate modulation of proinflammatory cytokine productions (55). The complex is resident in the nucleus in resting cells but migrates to the cytoplasm in response to stimuli (12). p38α lacks NES; however, it is transported to the cytoplasm in complex with MK2 that bears an NES. MK2-NES remains masked in its inactive form, phosphorylation (Thr334 in human) of NES by p38α unmasks it (12, 56, 57).

As mentioned earlier, MAPK pathways represent one of the most ancient eukaryotic signal transduction pathways throughout evolution. In *C. elegans* PMK3 and MAK2 are implicated in the axon regeneration process downstream of DLK1 (MAP3K) and MKK4 (MAP2K) (21, 23). The DLK1 pathway is negatively regulated by the E3 ubiquitin ligase RPM-1/Highwire/Phr in worm, fly and mice in order to stabilize the synaptic connections (18–20, 58). PMK3 and MAK2 were shown to interact by yeast two hybrid assay and in a co-immunoprecipitation assay performed in HEK293T cells (21). As expected PMK3 was able to phosphorylate MAK2 *in vitro.* We found that, PMK3 tends to form large oligomeric forms in solution that is catalytically inactive along with catalytically active low molecular weight fractions. PMK3 phosphorylates MAK2 at T173 and Thr285 in addition to other residues. A phospho-ablative double mutant of MAK2 (T173A T285A) failed to exert its role in axonal regeneration in nematode (15). Thr285 in MAK2 is equivalent to Thr334 in human MK2 located in the NES region, phosphorylation of which has been well established in the activation and nuclear export of MK2 and p38α:MK2 complex in human to exert their signaling roles. We found that the activated MAK2 phosphorylates its upstream kinase PMK3 at multiple locations including the TxY (T285-x-Y287) motif essential for PMK3 to attain its highest-activity state. This is a new insight which is unreported in the literature. Identification of doubly phosphorylated pT285pY287 bearing peptide exclusively in presence of MAK2 provokes us to propose this newly found feedback loop as a mechanism for fully activating PMK3 by its downstream kinase. However, allosteric autophosphorylation of PMK3 at T285 by virtue of its interaction with MAK2 through the docking site also contribute to its activation to a minor extent (Fig. 2C & 6C). Interaction of PMK3 with MAK2 through the canonical docking site on MAK2 enables trans-phosphorylation of the substrate kinase as expected (59) but does not enhance PMK3-autophosphorylation that requires catalytic activity of MAK2 as well (Fig. 2C & D, 6C) In addition, PMK3 specifically engages in such a feedback relationship with its downstream MAK2 but not with its upstream MKK4 (MAP2K) (Fig. 1F). Of note, MK2 (mammalian counterpart of MAK2) has been reported to participate in a feedback relationship with RIPK1 that downregulates RIPK1 activity (53, 54). We speculate that the feedback loop either may act as a fail-safe mechanism to activate PMK3 in a situation when MKK4 is either unavailable or inactivated; or it may help gain quicker and more robust response of this pathway as and when required (Fig. 2E). Since DLK1 signaling is used in multiple phenomena such as axon regeneration (23), degeneration (24, 60), and synaptic pruning (25) in response to nervous system injury, it might need a seamless mechanisms for robust activation and faithful transmission inside the cell. A critical dependence on two different p38 MAPK cascades involving DLK1 and MLK1 for axonal regeneration, indicates a robust cross-talk between parallel signaling kinases in neuronal injury response (61). Future course of studies will shed light on the scenario in the cellular context in presence and absence of MKK4.

Phosphorylation mediated regulation of kinases is a common theme in kinase biology. Distribution of phosphorylation sites on either kinase in this study suggested that the existing structural models with similar kinases from other organisms were inadequate to capture the possible modes of binding between PMK3 and MAK2 to enable such distribution of phosphorylation sites. In addition, both these kinases contained IDRs and showed theoretical propensity to form droplets. These observations led us to hypothesize that droplet/condensate formation through LLPS might be a plausible mechanism to accommodate the ensemble conformational space for this complex. In many cases, kinases have been found to depend upon scaffolding proteins to undergo phosphorylation by a trans-kinase or to attain a critical local concentration for self-phosphorylation. Activation of IKK-complex is a classic example of such mechanism wherein IKK-complex undergo activation by means of condensate formation through LLPS in the signaling context in a NEMO-dependent manner (62). Regulation of kinase function through LLPS-driven condensate formation have been reported in other cases as well (63–68). However, to the best of our knowledge, LLPS-mediated formation of kinase-condensate that relies upon the presence of a trans-phosphorylating kinase is not known. Here, we report that an aggregation-prone p38 MAPK (PMK3) is kept bio-available by its activating downstream kinase MAPKAPK (MAK2) by formation of condensates through LLPS. Furthermore, this LLPS-mediated condensate formation leads to productive association between the two kinases where a feedback phosphorylation relationship is functional. Our studies with transgenic *C. elegans* strongly suggest such functional association between these kinases in the organism, wherein MAK2 regulates the amount of PMK3 under normal and stress conditions. Interestingly, it has been shown recently that human MKK6:p38α complex is also highly dynamic in nature and the disordered N-terminal linker of the MAP2K (MKK6) plays critical role in determining specificity of the MAPK to be activated by phosphorylation (69). This report and our observations provoke us to speculate that MAPK activation across species might occur through a highly dynamical engagement of the MAPK with its upstream (MAP2K) or downstream (MAPKAP) kinases with ensuing conformational plasticity. Species specific regulatory nuances maybe required for fine tuning of the process.

Together, we propose a hitherto unreported regulatory mechanism for p38-MAP kinases wherein a critical signaling p38-MAP kinase is kept bio-available through physical interaction and condensate formation by its downstream substrate kinase that in turn activate the upstream kinase through a feedback phosphorylation loop (Fig. 6E). Interaction through the canonical docking-motif on MAK2 is necessary but not sufficient for the full activation of PMK3 through allosteric means. Full activation of PMK3 defined by double phosphorylation of the TxY-motif requires catalytic activity of MAK2. Future course of studies will shed light whether this is a rare, kinase-specific mechanism active in a particular organism or a more wide-spread aspect in kinase activation and regulation.

## Acknowledgements and funding sources

This study was primarily supported by DBT-Wellcome Trust India Alliance Intermediate Fellowship to SP (IA/I/15/1/501852) and Senior Fellowship to AGR (IA/S/22/1/506243), and intramural core funding from Bose Institute and BRIC-NBRC to SP and AGR, respectively. PR and AH thanks the University Grants Commission, Government of India, and SM thanks Bose Institute for graduate research fellowship. MN thanks Science and Engineering Research Board (GoI) for the National Post-doctoral fellowship (PDF/2022/001807). Authors thank Dr. Amitabha Majumdar (NCCS, Pune) for helpful discussion and the proteomics facility at CSIR-CCMB for mass spectrometric data collection.

## Author contribution

Conceptualization: SP; Experimental design: SP, AGR and PR; Investigation: PR, CM, PRT, AH, VB, SM (structural modeling) and MN (kinetic modeling); Data analysis: SP, PR, AGR, SRC, CM, PRT; Figure: SP, PR, CM, PRT, MN, SM; First draft: SP; Editing: PR, AGR, PRT, MN

## Supplementary Figures

**Figure S1:**
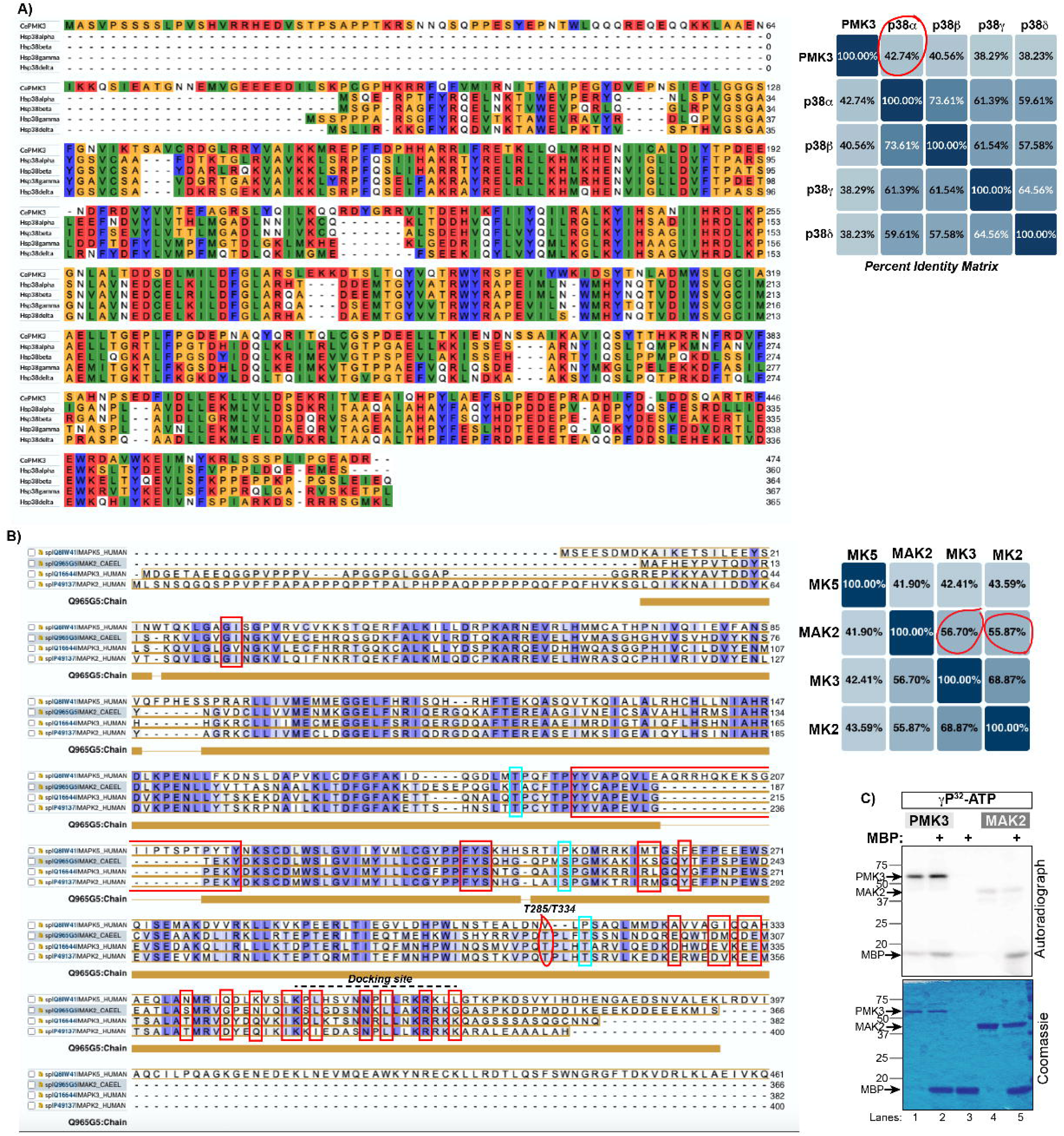
**(A)** Left panel: Multiple sequence alignment of C. elegans PMK3 (CePMK3) with the four human p38 (Hsp38) MAPK isoforms α, β, γ and δ. Right panel: The percent identity matrix obtained from the MSA of the p38 MAPK sequences. Darker shade in the box implies higher sequence identity. PMK3 shares highest sequence similarity with human p38α. **(B)** Left panel: Multiple sequence alignment (MSA) of MAK2 with the human MAPKAPKs MK2, MK3 and MK5. Right panel: The percent identity matrix shows that MAK2 shares more than 50% sequence similarity with MK2 and MK3. **(C)** *In vitro* radioactive kinase assay with purified, recombinant PMK3 and MAK2 using myelin basic protein (MBP) as a generic substrate (n=2).

**Figure S2:**
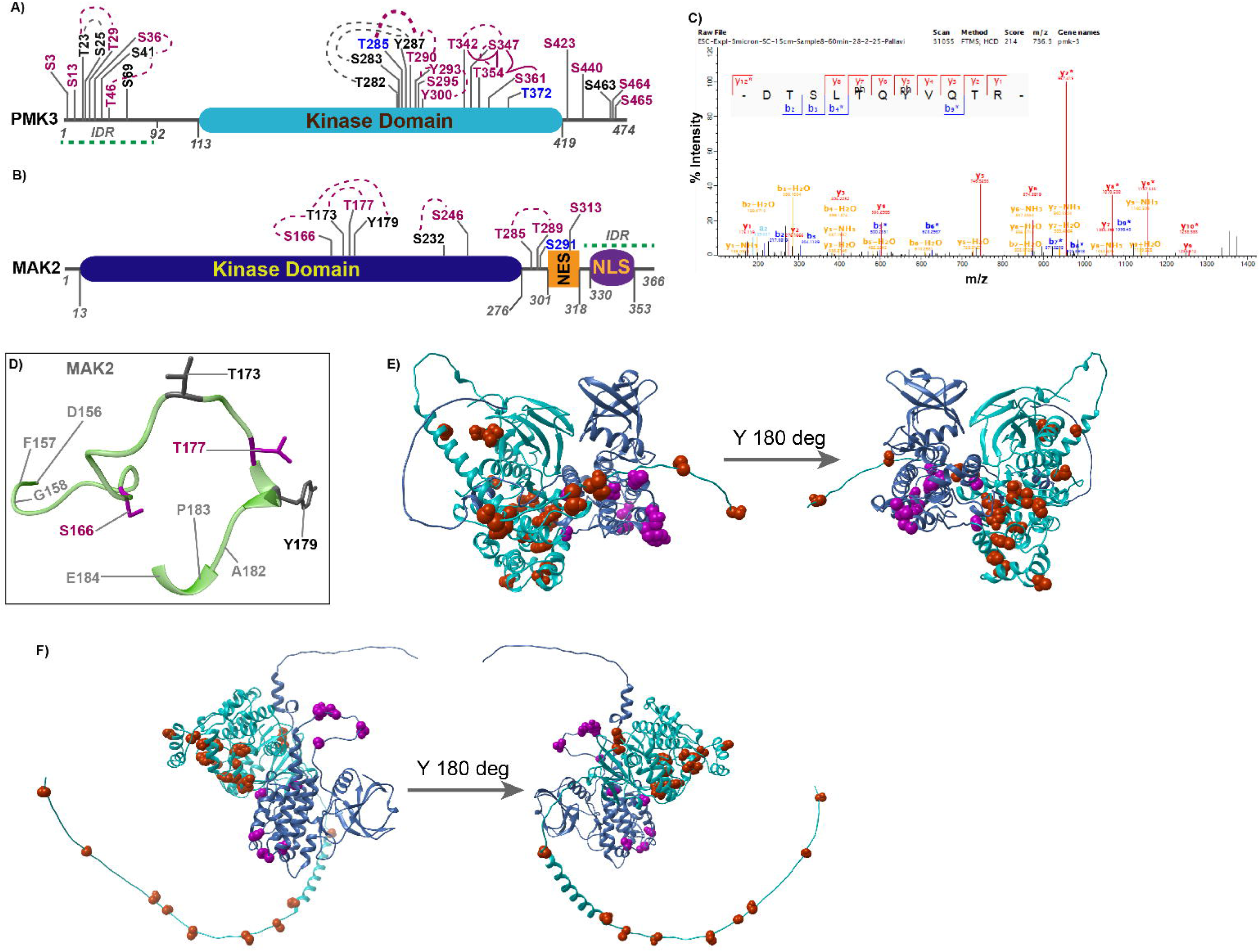
**(A-B)** Phosphorylation sites identified by LC-MS/MS experiments are shown on cartoon representations of PMK3 and MAK2 in *A* and *B*, respectively. Residues colored in blue were exclusively identified in individual kinases when the other kinase was absent, while those in purple were absent in the sample containing both kinases, and those in dark gray were identified in both types of samples. Residues connected with lines signify identification of those residues in doubly (dotted lines) or triply (solid lines) phosphorylated peptides. **(C)** MS/MS spectrum of a PMK3-peptide containing doubly phosphorylated TxY motif (DTSLpTQpYVQTR). **(D)** Ribbon representation of the activation segment of MAK2 from DFG- to APE-motif. Phosphorylation sites identified in this study are shown in sticks where in purple shows residues phosphorylated only in presence of PMK3 while those in dark gray are found both in presence and absence of PMK3. **(E)** Ribbon representation of the PMK3(cyan):MAK2(blue) complex obtained by superimposing on a p38α:MK2 (pdb id: 2oza). Phosphorylated residues identified by mass-spectrometric analyses have been highlighted (orange and purple) on both the kinases. **(F)** Ribbon representation of the AlphaFold2-predicted PMK3(cyan):MAK2(blue) complex. Phosphorylated residues identified by mass-spectrometric analyses have been highlighted (orange and purple) on those kinases.

**Figure S3:**
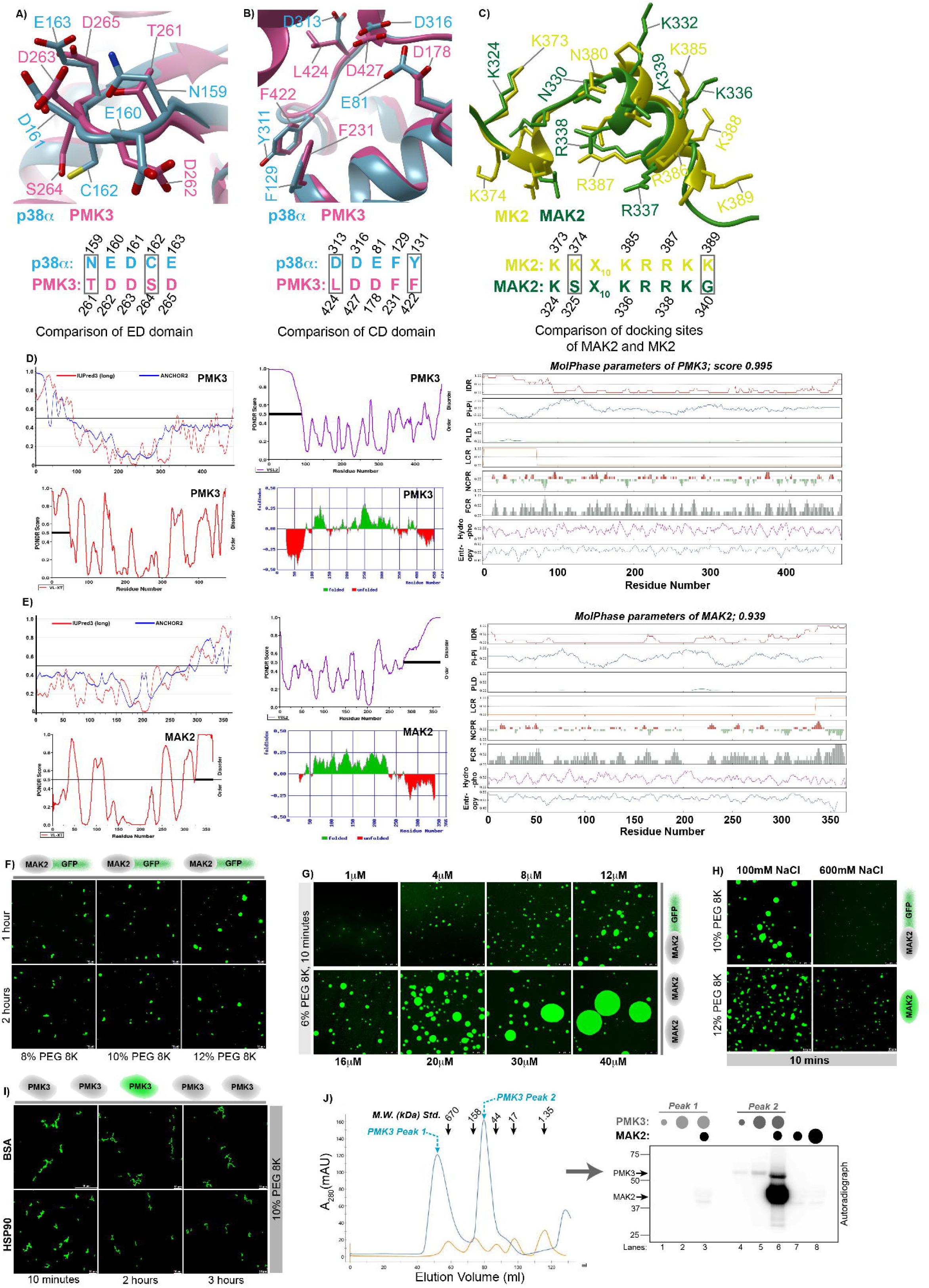
**(A-B)** Superimposition of the ED and CD domains of PMK3 and human p38α. The respective sequence alignments have been shown in the lower panel. Changes in the nature of amino acids have been highlighted in boxes. **(C)** Superimposition of the docking sites of MAK2 and MK2 notable changes are boxed. **(D-E)** IUPred3, PONDR, FoldIndex and MolPhase webserver-based predictions of PMK3 (D) and MAK2 (E) primary sequences. **(F)** A MAK2 construct with a GFP tag at its C-terminus (MAK2-GFP) could also successfully form these condensates through LLPS at the different PEG concentrations used. Images are representative of three independent experiments. Scale bar is 10μm. **(G)** Concentration dependent phase separation of MAK2-GFP in presence of 6% PEG 8K. MAK2-GFP was mixed with 6% PEG at different protein concentrations ranging from 1 to 40 μM and imaged under the microscope to monitor droplet formation. Scale bar is 8μm. **(H)** F-MAK2 and MAK2-GFP were subjected to phase separation in presence of increased amount of NaCl, wherein the droplet formation was significantly inhibited in both the cases at 600mM NaCl. Images are representative of two independent experiments. Scale bar is 8μm (upper panel) and 10μm (lower panel) respectively. **(I)** PMK3 forms aggregates even in the presence of chaperone Hsp90 or a non-specific protein, BSA. Scale bar is 10μm. **(J)** Left panel: Comparison of the size exclusion profiles of 6X His-PMK3 and a molecular weight standard resolved on a preparative gel filtration column. The absorbance at 280nm has been plotted against the elution volume. The larger molecular weight fractions constitute Peak 1 and the lower molecular weight fractions constitute Peak2. Right panel: *In vitro* radioactive kinase assay with purified, recombinant PMK3 and MAK2. PMK3 from two peaks in SEC was used for the assay in different reactions.

**Figure S4:**
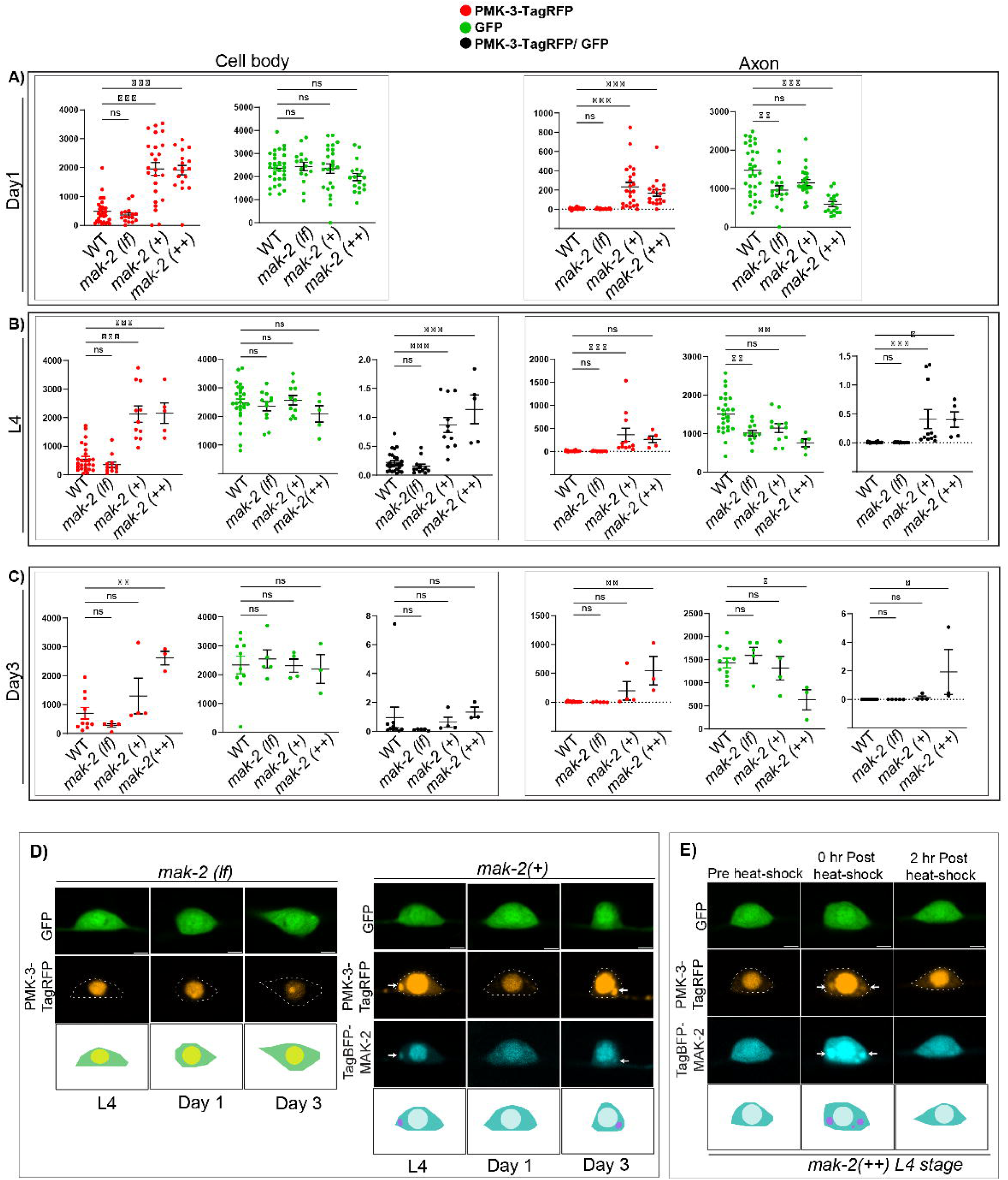
**(A)** Comparison of mean intensity of PMK3-TagRFP (Red-scatter plot) and diffusible GFP (Green-scatter plot) in region of interest in cell body and PLM axon in different genetic backgrounds at Day1 adult stage. **(B)** Comparison of mean intensity of PMK3-TagRFP, diffusible GFP and their ratio in region of interest in cell body and PLM axon in different genetic backgrounds at L4 stage. **(C)** Comparison of mean intensity of PMK3-TagRFP, diffusible GFP and their ratio in region of interest in cell body and PLM axon in different genetic backgrounds at Day3 adult stage. **(D)** High-resolution images of the cell body region of PLM neuron in mak-2(ok2394) background and in transgenic extrachromosomal array (shrEx526) co-expressing Pmec4-PMK3-TagRFP and Pmec4-TagBFP-MAK2 in 1:1 ratio (Scale bar - 2 μm). **(E)** High-resolution images of the PLM soma before and after heat shock in the transgenic line (shrEx527) co-expressing Pmec4-PMK3-TagRFP and Pmec4-TagBFP-MAK2 in 1:6 ratio.

**Figure S5:**
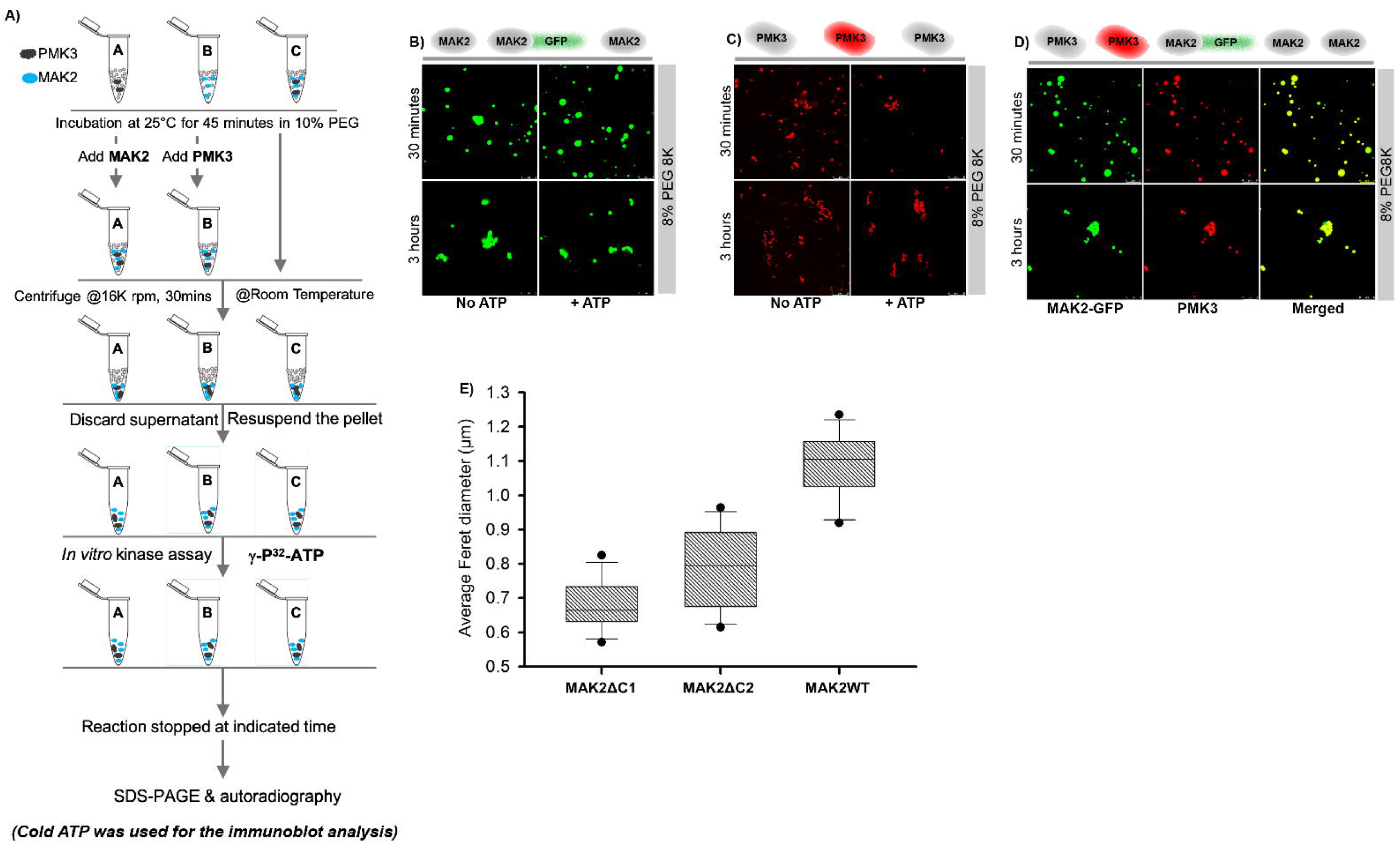
**(A)** Outline of the reaction schemes followed for the *in vitro* LLPS and subsequent kinase assays. **(B-C)** Condensate formation of MAK2 (B) and aggregation of PMK3 (C) in presence and absence of ATP. **(D)** Condensate formation of the PMK3:MAK2 complex through LLPS in absence of ATP. **(E)** Decrease in droplet size with the shortening of the C-terminal IDR of MAK2.

