## Supplementary material for "Activation and regulation of a p38-MAPK by its downstream MAPKAP kinase through feedback phosphorylation and LLPS-driven condensate formation": Materials and methods

***Materials***

All the chemicals used in this study were of molecular biology or analytical grade.  Growth media included LB broth (#M1245, HiMedia), LB agar (#M1151, HiMedia), SOC broth (#M1379, HiMedia) and Sf 900^TM^ III SFM (1X) media (#12658027, Gibco™, ThermoFisher Scientific). The ‘HF’ versions of all restriction enzymes were purchased from New England Biolabs. Other enzymes and related reagents included Calf Intestinal Alkaline Phosphatase (CIAP) (#18009019, ThermoFisher Scientific), TaKaRa Taq™ DNA Polymerase (#R001C, Takara Bio), Q5^®^ High-Fidelity DNA Polymerase (#M0491, NEB), Pfu DNA polymerase (#600382, Agilent Technologies), dNTP mixture (# 4030, Takara Bio), T4 DNA ligase (#EL0011, ThermoFisher Scientific), 10X T4 DNA ligase buffer (#B69, ThermoFisher Scientific). Antibiotics used were Carbenicillin (#C-103-5, GoldBio), Kanamycin (#K-120-25, GoldBio), Chloramphenicol (#C-105-25, GoldBio), Tetracycline (#103011, MP Biomedicals), Gentamicin (#15750078, ThermoFisher Scientific).

Antibodies and detection reagents included SuperSignal^TM^ West Pico PLUS Chemiluminescent Substrate (#34580, ThermoFisher Scientific), Clarity Western ECL Substrate (#1705061, Bio-Rad), purified Mouse anti-phosphotyrosine antibody (#610000, BD Biosciences), HRP-conjugated Goat anti-Rabbit IgG (H+L) (#31460, ThermoFisher Scientific) and Goat anti-Mouse IgG (H+L) Secondary Antibody (#31430, ThermoFisher Scientific).

Details of other reagents: Ni-NTA agarose (#30230, Qiagen), Agarose (#16500500, ThermoFisher Scientific), Lysozyme (#L6876, Merck), Bovine serum albumin (A-420-500, GoldBio), Protease inhibitor cocktail (#P8849, Merck), Adenosine-5’-triphosphate (ATP) (#A-081-25, GoldBio), Myelin basic protein (MBP) (#M1891, Merck), Polyethylene glycol 8000 (PEG 8000) (#V3011, Promega), Fluorescein-5-maleimide (#62245, ThermoFisher Scientific), Alexa Fluor 594 NHS ester (#A20004, ThermoFisher Scientific), Gel filtration standard (#1511901, Bio-Rad), HiTrapQ column (#17115301, Cytiva), Superdex 200 column (#28990944 or #90100137, Cytiva).

***STRING network analysis***

Multiprotein STRING search was performed on the STRING (version 12.0) database website using mak-2 and pmk-3 as input (1). For the full STRING network analysis, network edges were chosen to represent different types of interaction evidence, high confidence interactions with interaction scores ≥0.7 were selected and five or less interactors are displayed. Same parameter values were used for the physical subnetwork analysis on the same platform with these two proteins.

***Prediction of intrinsically disordered regions (IDRs) and droplet-promoting regions***

The amino acid sequences of *C. elegans* PMK3 (UniProt ID: O44514) and MAK2 (UniProt ID: Q965G5) were retrieved from the UniProt Knowledgebase (UniProtKB) in FASTA format. These sequences were used as inputs for the subsequent disorder and phase-separation propensity analyses. To identify intrinsically disordered regions, multiple web-based predictors were employed to ensure robustness of prediction. These included IUPred3 (<https://iupred3.elte.hu/>), PONDR® (<https://www.pondr.com/>) and FoldIndex© (<https://fold.proteopedia.org/cgi-bin/findex>). Regions consistently predicted as disordered by these tools were annotated as IDRs in our study. To assess the potential of IDRs to promote biomolecular condensation or phase separation, two specialized tools were employed: MolPhase (<https://molphase.sbs.ntu.edu.sg/>) and FuzDrop (<https://fuzdrop.bio.unipd.it/predictor>).

***Prediction of structural models of PMK3:MAK2 complex***

Two approaches were adopted to generate structural models of PMK3:MAK2 complex. In one approach, AlphaFold-predicted individual kinase structures were superimposed on the respective chains in the X-ray crystal structure of human p38α and MK2. Alternatively, AlphaFold Multimer v3 pipeline (2) along with ColabFold (3) was used to predict the PMK3-MAK2 complex structure. MMseqs2 was integrated into the pipeline for multiple sequence alignment (4) . Protein sequences were given as input in the form of multi-FASTA files containing one entry per chain. MMseqs2 was used to generate multiple sequence alignment searching against Uniprot database and HHsearch was used to retrieve structural templates (5). Structural predictions were obtained using the multimer model. Recycles per run was set to 3 and all five multimer models were then ranked based on interface predicted TM-score (ipTM) and mean predicted local distance difference test (pLDDT) confidence values (6). UCSF Chimera X and/or UCSF Chimera were used for model visualization and interface analyses in this study (7).

***Cloning of full-length and truncated expression constructs for recombinant protein over-expression and purification***

For recombinant protein production, the full-length coding sequence of C. elegans PMK3 protein was amplified by PCR using Q5 high-fidelity DNA polymerase (New England Biolabs) and subsequently subcloned into a modified pET24d-HisTEV expression vector between the EcoRI and XhoI restriction sites. The cloning workflow comprised amplification of the target insert, restriction digestion of both the PCR product and the vector, gel extraction and purification of the digested fragments, followed by ligation using T4 DNA ligase (Thermo Fisher Scientific). The ligation mixtures were transformed into chemically competent E. coli DH5α, and positive clones were validated through colony PCR, double digestion and confirmed by Sanger sequencing. The pET24d-HisTEV construct was engineered to include an **N-terminal 6×His tag** for affinity purification, followed by a **TEV protease cleavage site** to enable tag removal, if required. The pET24d vector system carries a T7 promoter for IPTG-inducible expression, along with a lacI gene and a kanamycin resistance marker, allowing for efficient selection and regulation in E. coli hosts. PMK3-deletion constructs (PMK3ΔN1 and PMK3ΔN2) were cloned into the same vector using EcoRI and XhoI sites following the protocol described above. Similarly, the full-length **MAK2 and truncated variants (MAK2ΔC1 and MAK2ΔC2)** coding sequences were cloned into the pET24d-HisTEV expression vector using BamHI and SalI restriction sites. For fluorescence-based detection under the microscope in downstream applications, a C-terminal GFP fusion construct of MAK2 was generated. For this purpose, a tri-molecular ligation strategy was employed involved two PCR amplified inserts MAK2 and GFP, and pET24d-HisTEV linearized with BamHI and SalI as the backbone. The MAK2 gene was amplified from the pET24-HisTEV:MAK2 construct using BamHI and SpeI flanking primers, and the **GFP** coding sequence was amplified from the pMAL325 plasmid using SpeI and SalI flanking primers. For heterologous expression in insect cells, the full-length **MKK4** gene was cloned within the BamHI and XhoI sites of the pFastBacHT B vector, which enables N-terminal His-tagged protein production using the baculovirus system. All restriction enzymes used were sourced from New England Biolabs unless indicated otherwise.

Oligonucleotide primers used in this study are listed in Table 2.

***Site-directed mutagenesis and cloning of kinase-dead variants***

To generate catalytically inactive or kinase dead variants for comparative biochemical analyses, missense point mutations (PMK3 K150M and MAK2 K42M) were introduced into the respective target genes via site-directed mutagenesis by overlap-extension PCR using complementary pair of primers containing desired base-mutation (Table 2) and *Pfu* Ultra DNA polymerase (Agilent Technologies). Post-PCR amplification, DpnI was added to the reaction mixture to selectively digest the methylated parental DNA template. The digested products were transformed into E. coli, and plasmids from positive clones were Sanger-sequenced to confirm the presence of the intended nucleotide substitutions.

***Culture growth conditions***

For recombinant protein expression, the plasmid constructs were transformed into E. coli Rosetta 2 (DE3) cells, a strain engineered to enhance expression of eukaryotic genes by supplying rare tRNAs. Transformed colonies were selected on LB-agar plates supplemented with 50 μg/ml kanamycin and 34 μg/ml chloramphenicol. A single colony was used to inoculate a primary overnight culture in Luria–Bertani (LB) medium supplemented with the same set of antibiotics, which in turn was used to seed a secondary culture containing 50 μg/ml kanamycin. The secondary culture was incubated at 37°C with shaking until the optical density at 600 nm (OD_600_) reached 0.5-0.6, at which point recombinant protein expression was induced by the addition of IPTG. Induced cultures were then incubated at 16°C for 18-20 hours with constant agitation to facilitate soluble protein expression. Cells were harvested by centrifugation at 4,500 x g for 20 minutes at 4°C and the resulting pellets were stored at -80°C until further use.

***Purification of recombinant kinases from E. coli***

For purification of 6XHis-tagged recombinant proteins, bacterial cell pellets were thawed on ice and resuspended in a lysis buffer composed of 50 mM Tris-HCl (pH 8.0), 200 mM NaCl, 10 mM Imidazole, 10% (v/v) Glycerol, 5 mM β-Mercaptoethanol (β-ME) and 1 mM PMSF. Lysozyme was added to the cell suspension at a final concentration of 0.2 mg/ml prior to sonication to facilitate mechanical lysis. Cell lysis was performed on ice using a probe-based ultrasonic processor (Cole-Parmer, USA) with the following settings: 15 s pulse on / 45 s off, 30% amplitude, for a total pulse duration of 15–30 minutes depending on the culture volume. Lysates were clarified by centrifugation at 16,000 x g for 1 hour at 4°C. The resulting supernatant containing the soluble His-tagged protein was collected and incubated with Ni-NTA agarose resin, pre-equilibrated with the lysis buffer. Binding was performed by rotating the mixture gently for 2 hours at 4°C. After binding, the unbound flow-through was collected and the resin was washed sequentially with 20 column volumes (CV) each of lysis buffer (Wash 1) and wash buffer (Wash 2) containing 25 mM Imidazole to remove non-specifically bound proteins. The target protein was subsequently eluted using an elution buffer containing 250 mM Imidazole. Eluted fractions were analysed by SDS-PAGE to verify the purity and molecular weight of the protein. To remove the N-terminal 6XHis tag from the purified protein, TEV protease cleavage was performed on the pooled elution fractions. Prior to cleavage, Imidazole was removed by overnight dialysis at 4°C against a buffer containing 50 mM Tris-HCl pH 8.0, 200 mM NaCl, 5% (v/v) Glycerol and 5 mM β-ME. Protein concentration was determined by measuring absorbance at 280 nm using a Multiskan SkyHigh Microplate Spectrophotometer (Thermo Fisher Scientific) and calculated using the Beer-Lambert equation:

*A= ε x c x l*

where A is the absorbance at 280 nm, ε is the molar extinction coefficient of the protein, c is the protein concentration and l is the path length.

TEV protease digestion was performed by adding TEV protease at a molar ratio of 1:10 (protease: substrate) to the dialyzed sample. Additionally, 2 mM EDTA and 1 mM DTT were added to the reaction mixture to ensure optimal cleavage conditions. The digestion was carried out on ice at 4°C for 12-16 hours. To remove His-tagged TEV protease and any undigested protein, the reaction mixture was passed through a small volume (200 μl) of Ni-NTA resin. The flow-through containing the cleaved, tag-free protein was collected and used as the input for the subsequent gel filtration step. In cases where TEV cleavage was not performed, the pooled Ni-NTA elution fractions were directly subjected to size exclusion chromatography.

Gel filtration was performed using an ÄKTA Pure FPLC system (Cytiva) equipped with a **HiLoad 16/600 Superdex 200 pg** column or a Superdex 200 Increase 10/300 GL column, depending on the protein amount. All chromatographic steps were carried out at 4°C to preserve protein stability. Protein samples (either TEV-digested or uncleaved) were subjected to centrifugation at 15,000 x g for 20 minutes at 4°C or alternatively filtered through a 0.22 μm syringe filter to remove any particulate matter. Samples were loaded onto the column pre-equilibrated with the gel filtration buffer consisting of 50 mM Tris-HCl pH 8.0, 100mM NaCl, 3% Glycerol, 5mM β-ME and 2mM EDTA. Elution was carried out isocratically at a constant flow rate of 1.0 ml/min and protein elution was monitored by absorbance at 280 nm. Eluted fractions were collected across the peak volume and the selected fractions were resolved by SDS-PAGE to assess purity and size. The SEC-purified fractions were further subjected to an additional round of purification to achieve higher purity of the recombinant proteins.

Anion exchange chromatography was carried out using a 5 ml HiTrap^TM^ Q HP column (Cytiva) connected to an ÄKTA Pure™ FPLC system (Cytiva). The column was equilibrated with Buffer A containing 50mM Tris-HCl pH 8.0, 50 mM NaCl, 3% (v/v) Glycerol, 5 mM β-ME and 2mM EDTA, prior to sample loading. Protein samples from pooled size exclusion fractions were either diluted or buffer-exchanged into Buffer A to reduce ionic strength and facilitate optimal binding to the anion exchange resin. Samples were loaded onto the column at a flow rate of 0.5–1.0 ml/min, followed by washing with 10 column volumes of Buffer A to remove unbound proteins. Bound proteins were eluted using a either a **stepwise or a linear salt gradient ranging from 50 mM to 2 M NaCl** over **30 column volumes**, achieved by mixing **Buffer A** and Buffer B in defined ratios. Fractions corresponding to the major elution peaks were collected and analysed by SDS-PAGE to confirm the presence and purity of the protein. Those containing the target protein were pooled and dialyzed against a buffer (50mM Tris-HCl pH 8.0, 100 mM NaCl, 3% (v/v) Glycerol, 5 mM β-ME and 2mM EDTA) to reduce the increased salt concentration introduced during elution. Fractions containing the desired protein were pooled and either used directly for downstream assays or concentrated at 4°C using centrifugal ultrafiltration devices (10 kDa or 30 kDa MWCO, Sartorius or Millipore). Final protein concentrations were determined spectrophotometrically, as described previously.

The specific expression conditions and purification steps for each kinase are summarized in Table 3.

***Protein purification from Sf9-baculovirus expression system***

The C. elegans His-tagged MKK4 protein was expressed in Spodoptera frugiperda (Sf9) insect cells. Recombinant bacmid generation, transfection into Sf9 cells and subsequent amplification of P1 to P3 viral stocks were performed according to the manufacturer's protocol (Thermo Scientific, MAN0000414) with some modifications. For large-scale expression, Sf9 cells maintained in suspension culture were infected with the high-titre P3 baculovirus at a density of 1 x 10^6^ cells/ml and incubated at 27°C for 60-72 hours under constant agitation (~125 rpm). Post-infection, the cultures were harvested by centrifugation at 1,200 x g for 30 minutes at 4°C. The cell pellets were washed once with ice-cold 1X TBS containing 2 mM EDTA and stored at -80°C until further use. Cell pellets were thawed on ice and resuspended in ice-cold lysis buffer comprising 50 mM Tris-HCl (pH 8.0), 200 mM NaCl, 10 mM Imidazole, 10% (v/v) Glycerol, 5 mM β-ME, 1 mM PMSF and 0.5X protease inhibitor cocktail (Sigma-Aldrich). Cell lysis was performed on ice by probe sonication with intermittent pulses (10 secs pulse on, 50 secs pulse off at 30% amplitude). The clarified lysate was applied to a Ni-NTA agarose affinity column pre-equilibrated with lysis buffer, and the bound His-tagged MK4 was purified as described in the previous section. To further enhance purity and remove aggregates or degradation products, the eluted fractions were pooled and subjected to size exclusion chromatography (SEC) using a HiLoad 16/600 Superdex 200 pg column (Cytiva) pre-equilibrated with SEC buffer consisting of 50 mM Tris-HCl (pH 8.0), 100 mM NaCl, 3% (v/v) Glycerol, 5 mM β-ME and 2 mM EDTA. Fractions corresponding to the observed peaks at 280 nm were analysed by SDS-PAGE, pooled and concentrated at 4°C using a centrifugal ultrafiltration device (30 kDa MWCO). The final protein preparation was stored at -80°C in small aliquots for downstream applications.

***In vitro radioactive kinase assays***

*In vitro* radioactive kinase assays were performed to assess the functional activity of the purified recombinant kinases and to investigate their mutual interactions. ɣ^32^P-ATP was used as the radiolabelled phosphate donor to monitor both auto- and trans-phosphorylation events. Reactions were assembled using the purified kinases, 200 μM non-radioactive (cold) ATP mixed with ɣ^32^P-ATP (hot ATP), in the presence of kinase assay buffer consisting of 20 mM HEPES (pH 7.7), 20 mM β-Glycerophosphate, 100 mM NaCl, 100 μM Sodium orthovanadate, 10 mM MgCl_2_, 10 mM Sodium fluoride, 1 mM PMSF and 2 mM DTT (added freshly). Where applicable, myelin basic protein (MBP) from Sigma-Aldrich was used as a generic kinase substrate. The protein concentration in each reaction was kept constant within a given experiment to ensure comparability and accurate interpretation of kinase activity. In general, kinase assay reactions were incubated at 25°C for varying time intervals and subsequently terminated by the addition of 4X SDS loading dye to achieve a final 1X concentration. Samples were then boiled for 5 minutes at 100°C and resolved by SDS-PAGE. To verify the amount of protein loaded in each reaction, gels were quickly stained with Coomassie Brilliant Blue and subsequently destained to visualize the protein bands. The stained gels were dried using a gel drying apparatus (Bio-Rad Model 583) and exposed to a phosphor screen (Cytiva) inside a cassette. Radioactive tracer signals were detected by scanning the phosphor screen using an Amersham Typhoon Imager (Cytiva).

***Identification of phosphorylation sites by LC-MS/MS***

To identify phosphorylation sites on the purified recombinant kinases, *in vitro* phosphorylation reactions were set up using 200 μM non-radioactive ATP in 1X kinase assay buffer supplemented with 2 mM DTT. Reactions were incubated at 25°C for 1 hour. Reactions were quenched by the addition of 20 mM EDTA. The final DTT concentration in each reaction was adjusted to 10 mM and the samples were incubated at 56°C for 45 minutes to reduce disulfide bonds. After cooling to room temperature, iodoacetamide (IAA) was added to a final concentration of 30 mM and the reaction was incubated for 1 hour in the dark to alkylate cysteine residues. Excess IAA was quenched by the addition of 5 mM DTT, followed by an additional 30-minute incubation at room temperature. Trypsin was reconstituted in 50 mM ammonium bicarbonate buffer and added to the samples at a 1:100 enzyme-to-substrate ratio (w/w). Trypsin was added in two steps: an initial addition was followed by incubation at 37°C for 1 hour, after which a second aliquot of trypsin was added and the digestion was continued overnight at 37°C. The reactions were terminated by acidifying the samples with formic acid to lower the pH below 3. The digested peptides were vacuum-dried using a SpeedVac (Thermo Scientific, Savant SPD111V), and phosphopeptides were selectively enriched using the High-Select Fe-NTA Phosphopeptide Enrichment Kit (ThermoFisher Scientific), following the manufacturer’s protocol.

Subsequently, the enriched peptides were desalted using Pierce C18 Tips (Thermo Scientific), eluted and resuspended in 5% (v/v) formic acid. Samples were briefly sonicated for 5 minutes and subjected to LC-MS/MS analysis. Peptides were analysed on an Orbitrap Exploris 240 mass spectrometer (Thermo Scientific) coupled to a nanoflow LC system (Easy-nLC II, Thermo Scientific). Samples were loaded onto a PepMap RSLC C18 nanocapillary reverse-phase column (75 μm x 25 cm, 2 μm particle size, 100 Å pore size) and separated using a 60-minute linear gradient of organic mobile phase consisting of 5% ACN with 0.1% formic acid (Buffer A) and 95% ACN with 0.1% formic acid (Buffer B). Raw MS data were processed using the MaxQuant computational proteomics platform (version 1.6.8) (8) and searched against UniProt amino acid sequences corresponding to the proteins of interest (UniProt IDs: Q965G5 and O44514) along with the *E. coli* proteome (UniProt ID: UP000002032). Phosphorylation of serine, threonine and tyrosine (STY) residues was specified as a variable modification in the search parameters. Phosphorylation sites with probability ≥0.85 that were detected at least twice (MS/MS count ≥2) have been reported. To compare percentage occurrence, only those phosphopeptides conforming to the above-mentioned criteria were considered and their MS/MS counts were summed up and was taken as total phosphopeptide occurrence. Percentage occurrence of each phosphorylation site was calculated accordingly. The MS-based proteomics data of these experiments have been deposited to the ProteomeXchange Consortium via the PRIDE partner repository with the dataset identifier PXD069356 (9).

***Kinetic modelling***

The detailed kinetic steps of phosphorylation processes are the following:

The Schematic of the phosphorelay kinetics of PMK3 (Y) and MAK2 (Z) mediated by MKK4 (X).

$X$ mediated phosphorylation of $Y$ ($Y\underset{\to}{X}Y_{p}$)

|  | $Y+X\begin{matrix} k_{2}^{+} \\ \rightleftarrows\\ k_{2}^{-} \end{matrix}YX\underset{\to}{k_{2}}Y_{p}+X$ | ( 1 ) |
| --- | --- | --- |

$Z_{p}$ mediated phosphorylation of $Y$ ($Y\underset{\to}{Z_{p}}Y_{p}$)

|  | $Y+Z_{p}\begin{matrix} {k'}_{2}^{+} \\ \rightleftarrows\\ {k'}_{2}^{-} \end{matrix}YZ_{p}\underset{\to}{k_{2}^{'}}Y_{p}+Z_{p}$ | ( 2 ) |
| --- | --- | --- |

$Y_{p}$ mediated phosphorylation of Z ($Z\underset{\to}{Y_{p}}Z_{p}$)

|  | $Z+Y_{p}\begin{matrix} k_{3}^{+} \\ \rightleftarrows\\ k_{3}^{-} \end{matrix}ZY_{p}\underset{\to}{k_{3}}Z_{p}+Y_{p}$ | ( 3 ) |
| --- | --- | --- |

$Y$ mediated phosphorylation of $Z$ (Z$\underset{\to}{Y}Z_{p}$)

|  | $Z+Y\begin{matrix} {k'}_{3}^{+} \\ \rightleftarrows\\ {k'}_{3}^{-} \end{matrix}ZY\underset{\to}{k_{3}^{'}}Z_{p}+Y$ | ( 4 ) |
| --- | --- | --- |

We considered a Michaelis-Menten kinetic model, where the phosphorylation takes place through an intermediate complex formation, which either produce the phosphorylated form of the kinase or revert back to the kinase form. The respective rate constants for each kinetic step are provided in the kinetic scheme (Eqs. (1-4)). We employ a coarse-grained approach for dephosphorylation processes ($Y_{p}\to Y$ and $Z_{p}\to Z$), assuming a direct conversion from the phosphorylated kinase back to the kinase form. This simplification is appropriate when the focus lies on capturing the overall effect of phosphorylation kinetics on the larger network. We, now, formulate the system of coupled ordinary differential equations (CODEs) that describe the temporal evolution of each component considering both phosphorylation and the simplified dephosphorylation processes.

|  | $\frac{d[Y]}{dt}=-k_{2}^{+}\left[ Y \right]\left[ X \right]+k_{2}^{-}\left[ YX \right]-k_{2}^{'+}\left[ Y \right]\left[ Z_{p} \right]+k_{2}^{'-}\left[ YZ_{p} \right]+k_{-2}\left[ Y_{p} \right]-k_{3}^{'+}\left[ Z \right]\left[ Y \right]+(k_{3}^{'}+k_{3}^{'-})[ZY]$ | ( 5 ) |
| --- | --- | --- |

|  | $\frac{d[Y_{p}]}{dt}=k_{2}\left[ YX \right]+k_{2}^{'}\left[ YZ_{p} \right]-k_{-2}\left[ Y_{p} \right]-k_{3}^{+}\left[ Z \right]\left[ Y_{p} \right]+(k_{3}+k_{3}^{-})[ZY_{p}]$ | ( 6 ) |
| --- | --- | --- |

|  | $\frac{d[Z]}{dt}=-k_{3}^{+}\left[ Z \right]\left[ Y_{p} \right]+k_{3}^{-}\left[ ZY_{p} \right]-k_{3}^{'+}\left[ Z \right]\left[ Y \right]+k_{3}^{'-}\left[ ZY \right]+k_{-3}\left[ Z_{p} \right]$ | ( 7 ) |
| --- | --- | --- |

|  | $\frac{d[Z_{p}]}{dt}=-k_{2}^{'+}\left[ Y \right]\left[ Z_{p} \right]+\left( k_{2}^{'}+k_{2}^{'-} \right)\left[ YZ_{p} \right]+k_{3}\left[ ZY_{p} \right]+k_{3}^{'}\left[ ZY \right]-k_{-3}\left[ Z_{p} \right]$ | ( 8 ) |
| --- | --- | --- |

|  | $\frac{d\left[ YX \right]}{dt}=k_{2}^{+}\left[ Y \right]\left[ X \right]-{(k}_{2}+k_{2}^{-})\left[ YX \right]$ | ( 9 ) |
| --- | --- | --- |

|  | $\frac{d\left[ YZ_{p} \right]}{dt}=k_{2}^{'+}\left[ Y \right]\left[ Z_{p} \right]-\left( k_{2}^{'}+k_{2}^{'-} \right)\left[ YZ_{p} \right]$ | ( 10 ) |
| --- | --- | --- |

|  | $\frac{d\left[ ZY_{p} \right]}{dt}=k_{3}^{+}\left[ Z \right]\left[ Y_{p} \right]-(k_{3}+k_{3}^{-})[ZY_{p}]$ | ( 11 ) |
| --- | --- | --- |

|  | $\frac{d\left[ ZY \right]}{dt}=k_{3}^{'+}\left[ Z \right]\left[ Y \right]-(k_{3}^{'}+k_{3}^{'-})[ZY]$ | ( 12 ) |
| --- | --- | --- |

In the above CODEs, the rate constants for dephosphorylation of $Y_{p}$ and $Z_{p}$ are denoted as $k_{-2}$ and $k_{-3}$, respectively. We define the Michaelis-Menten constants for each phosphorylation steps (Eqs. (1-4)) as,

|  | $K_{M}^{\left( 2 \right)}=\frac{k_{2}+k_{2}^{-}}{k_{2}^{+}}, K_{M}^{'\left( 2 \right)}=\frac{k_{2}^{'}+k_{2}^{'-}}{k_{2}^{'+}}, K_{M}^{\left( 3 \right)}=\frac{k_{3}+k_{3}^{-}}{k_{3}^{+}}, K_{M}^{'\left( 3 \right)}=\frac{k_{3}^{'}+k_{3}^{'-}}{k_{3}^{'+}}$ | ( 13 ) |
| --- | --- | --- |

For $X$ regulated phosphorylation of $Y$ (Eq. (1)), we define a dimensionless parameter $A$, called activity of the corresponding regulatory molecule which modulate the phosphorylation process of the downstream kinase. Specifically, for $X$ mediated phosphorylation of $Y$ (Eq. (1)), we define the activity of $X$ as $A_{X}=k_{2}/K_{M}^{(2)}$. Similarly, for $Y_{p}$ mediated phosphorylation of $Z$ (Eq. (3)) activity of $Y_{p}$ is defined as $A_{Y_{p}}=k_{3}/K_{M}^{(3)}$ and for $Y$ mediated phosphorylation of $Z$ (Eq. (4)) activity of $Y$ is defined as $A_{Y}=k_{3}^{'}/K_{M}^{'(3)}$. In the feedback loop where $Z_{p}$ induces phosphorylation of $Y$ (Eq. (2)), the activity of $Z_{p}$ is defined as $A_{Z_{p}}=k_{2}^{'}/K_{M}^{'(2)}$. The activity, $A_{Z_{p}}$, also represents feedback sensitivity and strength (FSS). A higher value of $A_{Z_{p}}$ indicates stronger and more sensitive feedback regulation. Specifically, a higher $k_{2}^{'}$ value increases the strength of feedback regulation, while a lower $K_{M}^{'(2)}$ value signifies higher sensitivity in the feedback regulation. In our model, we adjust the FSS ($A_{Z_{p}}$) to investigate the role of $Z_{p}$ feedback regulation in the dynamics of phosphorylation kinetics. The activities of the regulators $X, Y$, and $Y_{p}$ ($A_{X}$, $A_{Y}$, and $A_{Y_{p}}$, respectively) can also be described in similar fashion in terms of regulatory sensitivity and strength.

*Parameters used:*

Total concentration of $X$, $\left[ X \right]=5 \mu M$

Total concentration of $Y$, ${[Y]}_{\mathrm{Total}}=10 \mu M$

Total concentration of $Z$, ${[Z]}_{\mathrm{Total}}=10 \mu M$

Where,

$${[Y]}_{total}=\left[ Y \right]+\left[ Y_{p} \right]+\left[ YX \right]+\left[ YZ_{p} \right]+\left[ ZY_{p} \right]+[ZY]$$

$$\left[ Z \right]_{total}=\left[ Z \right]+\left[ Z_{p} \right]+\left[ YZ_{p} \right]+\left[ ZY_{p} \right]+\left[ ZY \right]$$

$$k_{2}^{+}=1.0 {\mu M}^{-1}s^{-1}, k_{2}^{-}=0.9 s^{-1}, k_{2}=0.1 s^{-1}, k_{-2}=0.3 s^{-1}$$

$$k_{3}^{+}=2.0 {\mu M}^{-1}s^{-1}, k_{3}^{-}=0.9 s^{-1}, k_{3}=0.1 s^{-1} , k_{-3}=0.3 s^{-1}$$

$$k_{3}^{'+}=0.1 {\mu M}^{-1}s^{-1}, k_{3}^{'-}=0.9 s^{-1}, k_{3}^{'}=0.1 s^{-1}$$

$$k_{2}^{'-}=1.0 s^{-1}, k_{2}^{'}=1.0 s^{-1}$$

Based on these parameter values, the Michaelis-Menten constants take the values $K_{M}^{\left( 2 \right)}=1 \mu M$, $K_{M}^{\left( 3 \right)}=0.5 \mu M$, and $K_{M}^{'\left( 3 \right)}=10 \mu M$. Similarly, the activity values are, $A_{X}=0.1$, $A_{Y}=0.01$, and $A_{Y_{p}}=0.2$.

We tune $k_{2}^{'+}$ so that $K_{M}^{'\left( 2 \right)}(=1/k_{2}^{'+})$ takes on different values allowing us to ultimately tune $A_{Z_{p}}$. This approach helps us explore how variations in FSS influence the phosphorylation kinetics.

***Fluorescent labelling of proteins***

Fluorescent labelling of purified recombinant kinases was performed to enable their detection during *in vitro* droplet formation assays for assessing liquid-liquid phase separation (LLPS). Labelling was carried out using either Fluorescein-5-Maleimide or Alexa Fluor 594 NHS Ester. For MAK2 and its C-terminal deletion truncates MAK2ΔC1 and MAK2ΔC2,labelling was performed using Fluorescein-5-Maleimide, which specifically reacts with accessible thiol groups. The purified proteins were buffer-exchanged into a labelling-compatible buffer containing 20 mM potassium phosphate (pH 7.2), 100 mM NaCl, 3% Glycerol, 2 mM EDTA and 50 μM TCEP. Fluorophore was added at a molar ratio of 8:1 (dye: protein) and the mixture was incubated on ice for 4-5 hours with intermittent mixing. The reaction was quenched by the addition of 5 mM DTT. Excess free dye was removed using a PD SpinTrap G-25 desalting column (Cytiva), followed by dialysis into a buffer composed of 50 mM Tris-HCl (pH 8.0), 100 mM NaCl, 3% Glycerol, 5 mM β-ME and 2 mM EDTA. For PMK3, labelling was performed with both Fluorescein-5-Maleimide and Alexa Fluor 594 NHS Ester. In the case of Alexa Fluor 594 labelling, the protein was first dialyzed against 20 mM potassium phosphate buffer (pH 8.0), 100 mM NaCl, 3% Glycerol, 5 mM β-ME and 2 mM EDTA. The dye was added to the protein at a molar ratio of 1:5 (protein: dye) and processed as described above. Labelled proteins were resolved by SDS-PAGE and scanned in an Amersham^TM^ Typhoon Imager (Cytiva) using appropriate filters to assess labelling efficiency and verify protein integrity. The labelled proteins were subsequently used to spike the corresponding unlabelled kinases at 10% of the total protein volume during the *vitro* droplet assays. This approach enabled fluorescent visualization of phase-separated condensates under a confocal microscope without significantly perturbing the overall behaviour of the protein assemblies.

***In vitro droplet formation assay***

*In vitro* liquid-liquid phase separation (LLPS) of the purified proteins was assessed in the presence or absence of a crowding agent under controlled buffer conditions. Reactions were assembled in a dilution buffer composed of 50 mM Tris-HCl (pH 8.0), 50 mM NaCl, 2% Glycerol, 5 mM β-ME and 2 mM EDTA. LLPS was induced by mixing the protein samples at various concentrations and conditions with polyethylene glycol (PEG-8000; Promega) at defined final concentrations (v/v). Samples were incubated at 25°C for different time intervals as specified in individual experiments. In these experiments, to enable visualization of protein condensates, the reaction mixtures were spiked with 10% fluorescently labelled protein — either general fluorophore-labelled kinase or MAK2-GFP for some assays involving MAK2. Phase separation was initially monitored by the appearance of sample turbidity and subsequently confirmed by imaging under a confocal microscope. For imaging, 12 μl of each reaction mixture was placed on a clean glass slide and gently overlaid with a coverslip. Fluorescence imaging of the labelled proteins within droplets was performed using either a Leica TCS SP8 confocal microscope (63X oil immersion objective) or a Leica Stellaris 5 laser scanning confocal microscope (100X oil immersion objective), both equipped with Las X (SuiteX) acquisition software. Further image processing was performed using ImageJ (v1.54g) software.

***Quantification of phase separated droplet size***

For comparison of the droplet sizes between the phase separated condensates of fluorescein labelled MAK2, MAK2ΔC1 and MAK2ΔC2, four representative images from each of three independent experiments were analysed per condition. The average Feret diameter of the droplets was measured using the ImageJ (v1.54g) software. Box plots for each data set was generated in SigmaPlot 12 for statistical comparison.

***Fluorescence recovery after photobleaching (FRAP)***

To assess the internal molecular dynamics and mobility of proteins within phase-separated condensates, fluorescence recovery after photobleaching (FRAP) was performed on MAK2-GFP droplets. FRAP experiments were conducted within 10-20 minutes of initiating the LLPS reactions to ensure analysis of freshly formed, spherical condensates. Experiments were carried out using the FRAP module integrated into the Leica Stellaris 5 laser scanning confocal microscope. Droplets were identified and imaged using the 488 nm excitation laser line for GFP, set to 100% laser power for bleaching. A high-magnification zoom was used to focus on individual droplets. Prior to photobleaching, three pre-bleach images were acquired to establish baseline fluorescence. A circular region of interest (ROI) within a selected droplet was then bleached by continuous laser exposure for 4 seconds. Recovery of fluorescence within the bleached ROI was monitored for a total duration of 90 seconds, with image acquisition every 10 seconds. The fluorescence intensity at each post-bleach time point was normalized to the pre-bleach baseline intensity to account for any photobleaching due to imaging. The resulting recovery curves were plotted to evaluate the extent and kinetics of fluorescence recovery, thereby inferring the dynamic exchange properties of the labelled proteins within the condensates.

***Immunoblotting using p-tyrosine antibodies***

To detect pan-tyrosine phosphorylation on PMK3 and MAK2, *in vitro* kinase assays were performed using 200 μM non-radioactive ATP in 1X kinase assay buffer. Reactions were incubated at 25°C for the indicated time intervals and terminated by the addition of SDS loading dye. Samples were boiled at 100°C for 5 minutes, resolved by SDS-PAGE and subsequently transferred onto PVDF membranes. Membranes were blocked for 1 hour at room temperature in 5% BSA prepared in 1X TBS-T, followed by overnight incubation at 4°C with anti-phosphotyrosine primary antibodies (BD Biosciences). After three washes with TBS-T (15 minutes each), membranes were incubated for 1 hour at room temperature with an HRP-conjugated anti-mouse IgG secondary antibody (Thermo Fisher Scientific) and visualized using enhanced chemiluminescence (ECL) reagents.

***C elegans transgenes and related molecular cloning***

To express *pmk-3* in the touch-neuronal circuit, an expression Gateway entry clone pCR8::*pmk-3*::TagRFP was made (10). *pmk-3* (1425bp) was amplified from pCR-8::PMK-3 (cDNA) (pNBRGWY28) using forward primer AGR1372 and reverse primer AGR1373. The vector backbone (3330bp) was amplified from pCR-8:: *ptrn-1*:: TagRFP (pNBR45) using forward primer AGR1374 (5' atggtgtctaagggcgaagagc 3') and reverse primer AGR608 (5' ggctccgaattcgccctt 3'). The two fragments were ligated using In-Fusion® HD kit. Further, pCR8::*pmk-3*::TagRFP was LR (Invitrogen) recombined with the P*mec-4* Gateway destination vector (pCZGY553) to generate P*mec-4*::*pmk-3*::TagRFP (6643bp).

Similarly, To express *mak-2* in the touch-neuronal circuit, P*mec-4*::TagBFP::*mak-2* (6409bp) was cloned (11). The three fragments were ligated using the In-Fusion® HD kit. The vector backbone (4507bp) was amplified from P*mec-4*::mScarlet (pNBRGWY54) using forward primer AGR648 and reverse primer AGR649. TagBFP (801bp) was amplified from plasmid #124343 (Addgene) using forward primer AGR1386 and reverse primer AGR1387. *mak-2* (1101bp) insert was amplified from the *mak-2* (C44C8.6a) c-DNA (yk2020f16 clone) of Yuji Kohara collection using forward primer AGR1388 and reverse primer AGR1389. Primer sequences are provided in Table 2.

The cloned constructs were micro-injected into the gonads of young adults (muIs32) with Pttx-3-RFP or Pmyo-2-mCherry as co-injection marker. F1s were single-selfed and lines with high-transmission efficiency were used for experiments. Detailed information on the transgenic *C. elegans* lines are furnished in Table 4.

***Imaging and quantification***

Worms were immobilized with 10 mM levamisole in M9 buffer on 5% agarose pad. Nikon Confocal Microscope (AXR) was used for imaging. 561nm, 488nm and 405nm excitation lasers were used with Minimal Crosstalk mode that allows Sequential acquisition of TagRFP, GFP and TagBFP. Z-stacked images were captured with 0.5 μm slice interval. Image-J (Fiji) was used to extract 0.5 μm single slice image of PLM soma.

*For florescence intensity comparison:*

Quality sequential acquisition with Minimal Crosstalk mode and 60X oil objective was used for imaging. Images were captured with 1 AU pinhole size at fixed (1024 x 512) pixel resolution such that PLM soma and around ~100 μm axon is imaged. Averaging of 2 and dwell time of 2 μS was used.

Images are z-projected with Image-J (Fiji) and a fixed dimension elliptical ROI (major axis=6 μm, minor axis=4 micron) is drawn over the PLM soma and ~60 μm line ROI of 5 pixel thickness is drawn over the axon. Mean intensity values of TagRFP and GFP is obtained which are background corrected and used for quantification.

*For PLM soma puncta analysis:*

Fast sequential acquisition with Minimal Crosstalk mode and 100X oil objective was used for imaging. Images were captured with 1.4 AU pinhole size at 1024 X 1024 pixel resolution. Averaging of 1 and dwell time of 2 μS was used.

Number of puncta in soma were counted by visual inspection only. 0.5 to 1 μm visibly big blobs are called puncta. In case of co-injection, if blobs seen in both channels, then called puncta. While comparing puncta phenotype, each genotype was imaged at same imaging parameters.

***Heat-shock experiments***

Worms were maintained at 20°C and were incubated at 34°C for 2 h at L4, Day1 and Day3 adult stage in separate experiments. To score condensate recovery after heat stress, worms were allowed to recover at 20 degrees for 2 hours and imaged.

**Table 2: Oligonucleotide primers used in this study**

| **Name** | **Sequence** |
| --- | --- |
| MKK4_BamHI_F | CGCGGATCCATGGTTCAAGAAGATGAC |
| MKK4_XhoI_R | CCGCTCGAGCTATCCTCTGTGGTCGA |
| PMK3_EcoRI_Fwd | CGGAATTCATGGCGTCAGTCCCAT |
| PMK3_XhoI_Rev | CCGCTCGAGTCAGCGATCTGCTTCTCC |
| MAK2_BamHI_F | CGCGGATCCATGGCTTTTCATGAGTAT |
| MAK2_SalI_R | ACGCGTCGACCTAGGAAATCATTTTTTC |
| PMK3_K150M_F | CGTCGTTATGTTGCAATAATGAAGATGCGAGAACCGTTC |
| PMK3_K150M_R | GAACGGTTCTCGCATCTTCATTATTGCAACATAACGACG |
| MAK2_K42M_F | GACAAATTCGCGCTAATGGTGCTCCGAGACACA |
| MAK2_K42M_R | TGTGTCTCGGAGCACCATTAGCGCGAATTTGTC |
| PMK3_45F_EcoRI | CGGAATTCAACACCTGGTTGCAACAA |
| PMK3_95F_EcoRI | CGGAATTCAAACGACGATTCCAATTT |
| ceMAK2-317endSalI | ACGCGTCGACTCAAACTCTCATTGAGGCCAATG |
| ceMAK2-341endSalI | ACGCGTCGACTCAACCTTTTCTCCTCTTCGCC |
| ceMAK2-SpeI-R | GACTAGTGGAAATCATTTTTTCCTCTTC |
| AMGFP-SpeI-F | GACTAGTGGATCCGGCGCACCTGGC |
| GFP-SalI-R | ACGCGTCGACTCACTTGTACAGCTCGTCCATG |
| AGR1372 | ctccgaattcgcccttATGGCGTCAGTCCCATCG |
| AGR1373 | CCCTGGAGAAGCAGATCGCatggtgtctaagggc |
| AGR648 | AAGGGCGAATTCGACCCAGC |
| AGR649 | AAGGGCGAATTCGGAGCC |
| AGR1386 | CAGGCTCCGAATTCGCCCTTatgtccgaactcatcaaggag |
| AGR1387 | GAAAAGCCATgttgagcttgtgtccgagc |
| AGR1388 | caagctcaacATGGCTTTTCATGAGTATCCTG |
| AGR1389 | GCTGGGTCGAATTCGCCCTTCTAGGAAATCATTTTTTC |

**Table 3: Expression conditions and purification procedures for individual kinases**

| **Construct** | **Expression host** | **Induction O.D. and temperature** | **IPTG concentration** | **Affinity Chromatography** | **Size exclusion chromatography** | **Anion exchange chromatography** |
| --- | --- | --- | --- | --- | --- | --- |
| pET24d-HisTEV: PMK3 | *E. coli* | O.D_600_ = 0.5-0.6, 16°C | 0.25 mM | Yes | Yes | Yes |
| pET24d-HisTEV: PMK3-KD | *E. coli* | O.D_600_ = 0.5-0.6, 16°C | 0.3 mM | Yes | Yes | Yes |
| pET24d-HisTEV: PMK3ΔN1 | *E. coli* | O.D_600_ = 0.5-0.6, 16°C | 0.25 mM | Yes | Yes | Yes |
| pET24d-HisTEV: MAK2 | *E. coli* | O.D_600_ = 0.5-0.6, 16°C | 0.2 mM | Yes | Yes | Yes |
| pET24d-HisTEV: MAK2-KD | *E. coli* | O.D_600_ = 0.5-0.6, 16°C | 0.25 mM | Yes | Yes |  |
| pET24d-HisTEV: MAK2ΔC1 | *E. coli* | O.D_600_ = 0.5-0.6, 16°C | 0.2 mM | Yes | Yes |  |
| pET24d-HisTEV: MAK2ΔC2 | *E. coli* | O.D_600_ = 0.5-0.6, 16°C | 0.2 mM | Yes | Yes |  |
| pET24d-HisTEV-GFP: MAK2 | *E. coli* | O.D_600_ = 0.5-0.6, 16°C | 0.2 mM | Yes | Yes |  |
| pFastBacHT B: MKK4 | Sf9 insect cells |  |  | Yes | Yes |  |

**Table 4: Mutant and transgenic strains used in this study**

| **Strain number** | **Genotype** | **Construction** |
| --- | --- | --- |
| NBR270 | muIs32 [Pmec-7-GFP + lin-15(+)] II | In lin-15(n765 ts), Pmec-7-GFP + lin-15(+) array was gamma-irradiated for chromosomal insertion. |
| NBR1134 | muIs32(Pmec-7-GFP)II; pmk-3(ok169)IV; shrEx521 [5ng Pmec-4-PMK-3-TagRFP + Pttx-3-RFP] LINE1(65%eff.) | 5ng Pmec-4-PMK-3-TagRFP + Pttx-3-RFP colony4 was injected in CZ8789 pmk-3 (ok169)IV; muIs32 (Pmec-7-GFP) II (65%eff.) |
| NBR1135 | muIs32(Pmec-7-GFP)II; pmk-3(ok169); shrEx522 [5ng Pmec-4-PMK-3-TagRFP + Pttx-3-RFP] LINE2(65%eff.) | 5ng Pmec-4-PMK-3-TagRFP + Pttx-3-RFP colony4 was injected in CZ8789 pmk-3 (ok169)IV; muIs32 (Pmec-7-GFP) II (65%eff.) |
| NBR1136 | muIs32(Pmec-7-GFP)II; shrEx522 [5ng Pmec-4-PMK-3-TagRFP + Pttx-3-RFP] LINE2(65%eff.) | NBR1135 was crossed with N2 males and F2s having Pttx-3-RFP were genotyped for homozygous wild-type pmk-3. |
| NBR1137 | shrEx523 [5ng Pmec-4-PMK-3-TagRFP + Pttx-3-RFP] LINE1(50%eff.) | 5ng Pmec-4-PMK-3-TagRFP + Pttx-3-RFP colony4 was injected in N2 (50%eff.) |
| NBR1138 | muIs32(Pmec-7-GFP)II; shrEx524 [5ng Pmec-4-TagBFP-MAK-2 + Pmyo-2-mCherry] LINE1(70%eff.) | 5ng Pmec-4-TagBFP-MAK-2 + Pmyo-2-mCherry colony3 was injected in muIs32(Pmec-7-GFP)II |
| NBR1139 | shrEx529[5ng Pmec-4-TagBFP-MAK-2 + Pmyo-2-mCherry] LINE1(68%eff.) | 5ng Pmec-4-TagBFP-MAK-2 + Pmyo-2-mCherry colony3 was injected in N2 (68%eff.) |
| NBR1140 | muIs32(Pmec-7-GFP)II; mak-2(ok2394)IV ; shrEx522 [5ng Pmec-4-PMK-3-TagRFP + Pttx-3-RFP] LINE2(65%eff.) | NBR1135 was crossed with mak-2(ok2394)IV males and F2s having Pttx-3-RFP were genotyped homozygous for wild-type pmk-3 and mak-2(ok2394). |
| NBR1141 | muIs32(Pmec-7-GFP)II; shrEx525 [5ng Pmec-4-PMK-3-TagRFP + 5ng Pmec-4-TagBFP-MAK-2 + Pttx-3-RFP] LINE1 (80%eff.) | 5ng Pmec-4-PMK-3-TagRFP + 5ng Pmec-4-TagBFP-MAK-2 + Pttx-3-RFP was injected in muIs32(Pmec-7-GFP)II (80%eff.) |
| NBR1142 | muIs32(Pmec-7-GFP)II; shrEx526 [5ng Pmec-4-PMK-3-TagRFP + 5ng Pmec-4-TagBFP-MAK-2 + Pttx-3-RFP] LINE2 (70%eff.) | 5ng Pmec-4-PMK-3-TagRFP + 5ng Pmec-4-TagBFP-MAK-2 + Pttx-3-RFP was injected in muIs32(Pmec-7-GFP)II (70%eff.) |
| NBR1143 | muIs32(Pmec-7-GFP)II; shrEx527 [5ng Pmec-4-PMK-3-TagRFP + 30ng Pmec-4-TagBFP-MAK-2 + Pttx-3-RFP] LINE1 (68%eff.) | 5ng Pmec-4-PMK-3-TagRFP + 30ng Pmec-4-TagBFP-MAK-2 + Pttx-3-RFP was injected in muIs32(Pmec-7-GFP)II (68%eff.) |
| NBR1144 | muIs32(Pmec-7-GFP)II; shrEx528 [5ng Pmec-4-PMK-3-TagRFP + 30ng Pmec-4-TagBFP-MAK-2 + Pttx-3-RFP] LINE2 (53%eff.) | 5ng Pmec-4-PMK-3-TagRFP + 30ng Pmec-4-TagBFP-MAK-2 + Pttx-3-RFP was injected in muIs32(Pmec-7-GFP)II (53%eff.) |
